# Modulating Cardiac Energetics in Cardio-Metabolic Syndromes: A mechanistic, hyperpolarized MR Trial of Ninerafaxstat Treatment

**DOI:** 10.1101/2024.04.24.591019

**Authors:** Moritz. J. Hundertmark, Adrienne. G. Siu, Violet Matthews, Andrew J. Lewis, James Grist, Jai Patel, Paul Chamberlin, Rizwan Sarwar, Arash Yavari, Hakim-Moulay Dehbi, Prashant Rao, Xu Shi, Shuning Zheng, Jeremy M. Robbins, Robert E. Gerszten, Michael P. Frenneaux, Ladislav Valkovič, Jack J.J.J Miller, Stefan Neubauer, Damian J. Tyler, Oliver J. Rider

## Abstract

**Background:** Type 2 diabetes (T2D) and obesity are key contributors for heart failure (HF)- development, especially for HF with a preserved ejection fraction (HFpEF). On a molecular basis, excessive use of fatty acids (FA) induces lipotoxicity which in turn promotes inflammation, reduces mitochondrial pyruvate dehydrogenase (PDH) activity and impairs myocardial energetics and -function. Harnessing in-vivo, real time measurement of cellular metabolism via hyperpolarized pyruvate MR, we aimed to assess the effects of ninerafaxstat, a selective FA oxidation inhibitor, on cardiac energetics, metabolism & diastolic function in patients with cardio-metabolic syndromes.

**Methods:** IMPROVE-DiCE was an open-label, mechanistic phase 2a trial. 21 participants received 200mg ninerafaxstat twice daily for four (n=5) or eight weeks (n=16). Myocardial energetics (phosphocreatine to adenosine triphosphate ratio, PCr/ATP), metabolism and function were assessed pre-& post-treatment using magnetic resonance imaging (MRI), ^31^P- and ^1^H-MR spectroscopy (MRS). We utilised hyperpolarized [1-^13^C]pyruvate MRS to assess in-vivo PDH-flux (n=9) and plasma metabolomics and proteomics to assess whole body metabolism.

**Results:** Patients presented with impaired PCr/ATP, (median 1.6 [IQR 1.4, 2.1]), myocardial steatosis (2.2 % [IQR 1.5, 3.2]) and LV diastolic dysfunction (peak circumferential diastolic strain rate 0.86/s [IQR 0.82, 1.06]) at baseline. Ninerafaxstat treatment improved myocardial energetics by 32% (*p*<0.01), reduced myocardial triglyceride content by 34% (*p*=0.03) and showed a trend towards improved PDH-flux (mean 45% increase, *p*=0.08). Diastolic function was significantly improved post-treatment (peak diastolic strain rate by 10%, peak LV filling rate by 11%, both *p*<0.05).

**Conclusions:** Metabolic modulation with ninerafaxstat significantly improved myocardial energetics, reduced myocardial steatosis and improved LV diastolic filling. Combining hyperpolarized MRS and metabolomics, is a powerful approach to examine the mechanism of action of novel metabolic modulators.

**REGISTRATION:** URL: https://clinicaltrials.gov; **Unique identifier:** NCT04826159

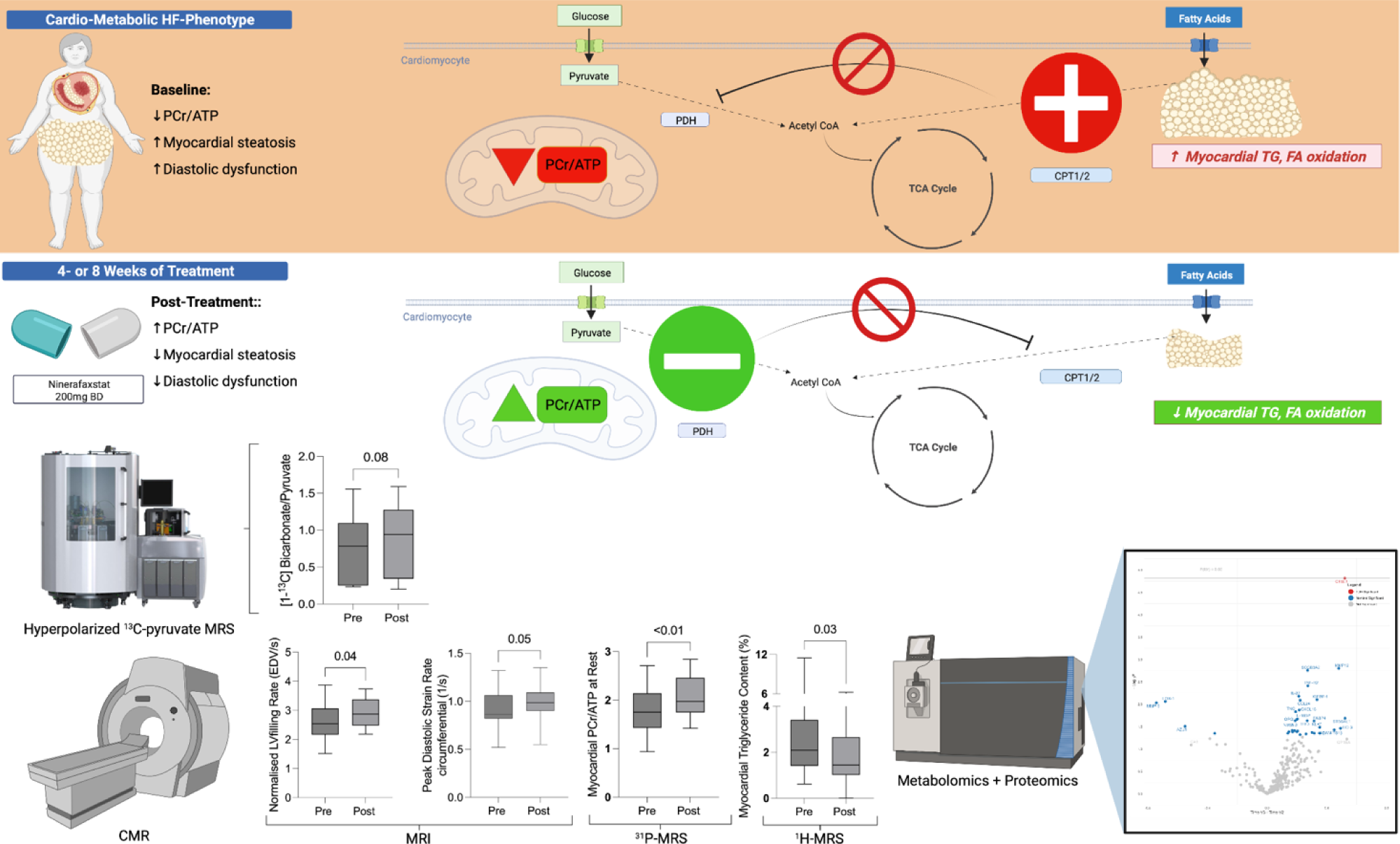

## Introduction

Type 2 diabetes (T2D) and obesity have long been misjudged as simple bystanders of heart failure (HF) development. However, lately this view has changed and both conditions are now seen as active modulators of HF-incidence, symptom severity and outcomes.^1^ Moreover, novel drugs targeting T2D and obesity in patients with HF with a preserved ejection fraction (HFpEF), with or without T2D, have emerged as important treatments.^2^ As the majority of patients suffer from both conditions and obesity begets T2D, concomitant existence of both disorders has been termed ‘diabesity’. Diabesity is associated with significant myocardial metabolic alterations linked to both structural and functional pathological changes, including left ventricular hypertrophy, diastolic and systolic dysfunction.^3^ The phenotype these patients typically present has been described as ‘cardio-metabolic syndrome’, which frequently represents an asymptomatic HF state (stage A or B) often progressing to clinically overt HF (stage C/D), which is then termed HFpEF.^4,5^ Diabesity is a common denominator for a majority of patients with HFpEF and is linked to a higher symptom burden and worse outcomes compared to non-diabese patients and thus, of significant importance to cardiologists.^6,7^ In addition, there is a substantial unmet medical need for novel therapies treating diabesity-related HF and preventing progression of asymptomatic cardio-metabolic syndromes to clinically overt HF.^8^

The heart is the highest energy consumer (adenosine triphosphate (ATP) per gram of tissue) in the body and thus, requires unmitigated energy supply with a complete turnover of its ATP pool every ∼10 seconds.^9^ As such, any mismatch between ATP-provision and demand leads to an energetic deficit and cardiac dysfunction, exacerbated further in the context of increased workload.^10^ Under physiological conditions, the healthy heart is metabolically flexible and able to dynamically use a range of carbon-based substrates to generate ATP. Free fatty acids (FFA) and glucose, as well as contributions from lactate, ketone bodies and several amino acids ensure close coupling of mitochondrial ATP production with the rate of ATP-use across a range of states such as feeding, fasting and exercise.^11^

Conversely, diabesity increases levels of circulating FFA and glucose ^12^ with an over-reliance on FFA to generate ATP, in the context of progressive insulin resistance.^13^ This results in a loss of metabolic flexibility ^14^, lipotoxicity and inhibition of pyruvate dehydrogenase (PDH) resulting in uncoupling of glycolysis from glucose oxidation.^6^ Over time, this ‘metabolic remodelling’ increases myocardial oxygen consumption, oxidative stress and inflammation but also reduces cardiac efficiency.^15^ Consequently, diabesity per se is associated with a lower phosphocreatine to ATP ratio (PCr/ATP, a sensitive marker of energetic status) evident early in the disease process.^16,17^ Hence, the diabetic heart displays reduced PDH flux^18^, impaired myocardial energetics^19^, cardiac steatosis^20^, left ventricular (LV) hypertrophy, and LV diastolic and/or systolic dysfunction.^21^.

Ninerafaxstat was designed to shift myocardial substrate utilisation in favour of glucose oxidation through partial FAO-inhibition (pFOX) and use of a precursor in the synthesis of NAD^+^ to enhance the cellular NAD^+^ pool. These effects aim to restore substrate flexibility and improve cardiac energetics, metabolism and function. In rodents, ninerafaxstat markedly increases myocardial glucose uptake and exerts protective effects in pre-clinical models of ischemia-reperfusion injury and pressure overload.^22^ Very recently, treatment with ninerafaxstat in patients with non-obstructive HCM was shown to be safe and effective, resulting in improved exercise performance.^23^

In this trial, we sought to examine the effects of ninerafaxstat on myocardial function, energetics and metabolism in patients with cardio-metabolic syndromes, using state of the art cardiovascular magnetic resonance (CMR) tri-nuclear (^31^P, ^1^H and ^13^C) MR-spectroscopy (MRS) to evaluate myocardial energetics, steatosis and PDH flux, respectively.

## Methods

IMPROVE-DiCE was a single-centre, open-label, proof-of-mechanism phase 2a trial designed to assess the effects of ninerafaxstat on myocardial energetics, metabolism, and function in participants with cardio-metabolic syndrome. The trial was registered (EudraCT Number: <u>2020-003280-26</u>, ClinicalTrials.gov Identifier: <u>NCT04826159</u>) and sponsored by Imbria Pharmaceuticals, Inc. Ethical approvals for the study were granted by the Medicines and Healthcare Products Regulatory Agency (MHRA) and National Research Ethics Service (Research Ethics Committee ref 20/LO/<u>1120</u>). The trial was conducted in accordance with the principles of the Declaration of Helsinki and the EU Clinical Trial Directive. All participants provided written informed consent prior to any investigations. Study visits took place at the Oxford Centre for Clinical Magnetic Resonance at the John Radcliffe Hospital, Oxford, United Kingdom.

### Participants and procedures

The population enrolled were adults aged 18-75 years old with T2D (HbA_1c_ ≥ 6.5 %), a body mass index (BMI) ≥ 30-40 kg/m^2^ and preserved left ventricular ejection fraction (LVEF ≥ 50 %). Exclusion criteria included treatment with insulin and/or sodium-glucose cotransporter-2 inhibitor (SGLT2i), more than moderate renal impairment (defined as an estimated glomerular filtration rate < 60 mL/min/1.73m^2^), NYHA III or IV heart failure, any change in oral antidiabetic therapy in the last 3 months and any contraindications to MR-scanning. A detailed list of in- and exclusion criteria can be found in *Table S1* in the *Data Supplement*.

A study flow chart is shown in *Figure 1*. The baseline visit included fasting blood sampling, cardiac MRI, resting and dobutamine ^31^P-MRS, as well as ^1^H- and [1-^13^C]pyruvate hyperpolarized MRS. Following baseline assessments, participants received ninerafaxstat administered orally at 200 mg BID on the day of their baseline visit, continuing for the duration of the respective treatment interval. The first 5 participants received ninerafaxstat for 4 weeks, with the subsequent 16 participants receiving ninerafaxstat for 8 weeks.

**Figure 1:**
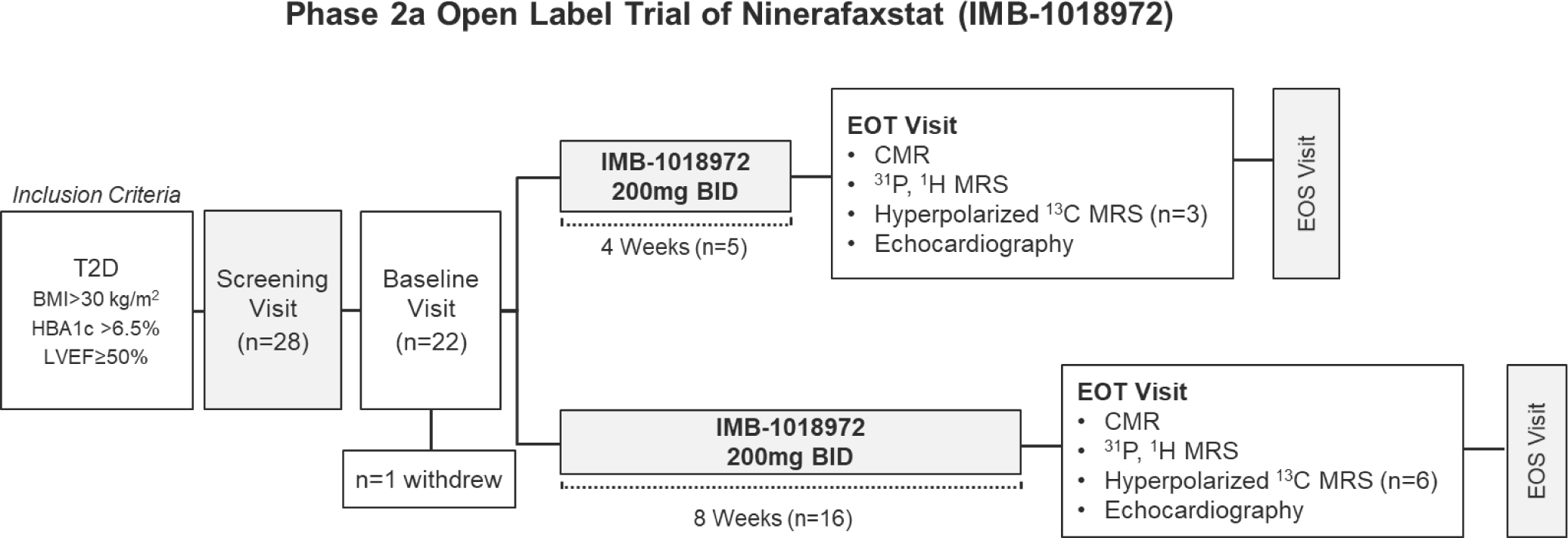
IMPROVE-DiCE patient flow and visit schedule. A schematic overview of the study design and visit schedule. ^1^H-MRS = proton magnetic resonance spectroscopy, ^31^P-MRS = phosphorus magnetic resonance spectroscopy, BID = twice daily, BMI = Body Mass Index, CMR = cardiovascular magnetic resonance, EOS = end of study, EOT = end of treatment, LVEF = left ventricular ejection fraction, T2D = type 2 diabetes.

### Trial oversight

Medpace, as the clinical research organisation, coordinated and monitored the study including safety monitoring and data management. A web-based electronic case record form was used. Adverse event reporting was performed by the study team and independently reviewed by Medpace. Initiation, routine monitoring, and closeout visit were performed on site.

### Study endpoints

The primary outcome of the study was the change from baseline to end-of-treatment in myocardial PCr/ATP ratio, a marker of cardiac energetic reserve, measured by ^31^P-MRS. Secondary outcomes included changes from baseline to end of treatment in cardiac systolic and diastolic function, systolic augmentation to dobutamine, and an evaluation of the safety and tolerability of repeat oral doses. Additionally, we sought to investigate the impact of ninerafaxstat on myocardial PDH-flux inferred by the [1-^13^C]bicarbonate to [1-^13^C]pyruvate ratio measured by hyperpolarized [1- ^13^C]pyruvate MRS in a subset of 9 participants.

### Exploratory outcomes

These included changes in: myocardial steatosis, myocardial enzymatic fluxes reflecting the metabolic fate of pyruvate, including the lactate dehydrogenase (LDH) and alanine aminotransferase (ALAT) reactions measured by hyperpolarized [1-^13^C]pyruvate MRS, systemic insulin sensitivity (Homeostasis Model Assessment [HOMA] and quantitative insulin sensitivity check index [QUICKI]), lipid profile and circulating N-terminal pro b-type natriuretic peptide (NT-proBNP) and high-sensitivity cardiac troponin (hs-cTn). Furthermore, we performed untargeted analyses of plasma proteomic and metabolomic markers pre- and post-treatment with ninerafaxstat in the entire trial cohort.

### Anthropometric and biochemical assessment

Height, weight and blood pressure were recorded at baseline and post-treatment. Following overnight fasting, participant’s venous blood samples were drawn and biomarkers analysed according to standardised protocols. Fasting insulin resistance was represented by HOMA-IR ((glucose (mmol/l) x insulin (pmol/l))/135).

### Cardiac magnetic resonance imaging and spectroscopy

A brief overview of techniques is described below with additional methodological details provided in the *Data Supplement*.

### MR imaging

Using balanced steady-state free precession cine imaging, electrocardiogram (ECG) gated long- and short axis images of the left ventricle (LV) were acquired.

Image analysis for biventricular indices was performed offline in accordance with Society for Cardiovascular Magnetic Resonance (SCMR) guidelines^24^, using cvi42 post-processing software (version 5.10.1, Circle Cardiovascular Imaging Inc., Calgary, Canada).

### 31P-MRS to assess myocardial energetics

^31^P-MRS was performed on a 3-Tesla MR scanner (MAGNETOM Trio, Siemens Healthineers, Erlangen, Germany). In brief, participants were positioned prone over the center of a 3-element dual-tuned ^1^H/^31^P surface coil (Siemens Medical, Erlangen, Germany) in the MR-isocentre. A non-gated 3D acquisition-weighted ultra-short echo time (UTE) chemical shift imaging sequence was used with saturation bands placed over liver and skeletal muscle, as previously described.^25^ All spectra were analysed using a semi-automated fitting of data utilising the OXSA toolbox,^26^ i.e. a MATLAB implementation of the AMARES fitting routine performed by an experienced operator (MH, 5 years CMR experience). Further details concerning data analysis are provided in the *Data Supplement*.

### Hyperpolarized [1-^13^C]pyruvate spectroscopy

A General Electric SpinLab system (GE Healthcare, Chicago, USA) was used for the process of Dynamic Nuclear Polarisation as described previously.^18^. Intravenous injection of the hyperpolarized [1-^13^C]pyruvate was undertaken at a dose of 0.4 mL/kg and at a rate of 5 mL per second via a power injector (MEDRAD, Bayer). ^18^ Further details regarding spectral analysis are provided in the *Data Supplement*.

### 1H-MRS to assess myocardial steatosis

All ^1^H-MR assessments were performed on a 3.0 Tesla MR system (Siemens, Germany) as previously described.^27,28^ In brief, end-expiration, ECG-triggered spectra were obtained from the mid inter-ventricular septum avoiding vascular structures. Spectra were acquired with and without water suppression to measure MTG-content. Spectra were analysed using Matlab and the AMARES algorithm within the OXSA toolbox. MTG-content was calculated as a percentage (signal amplitude of lipid/signal amplitude of water) ×100.

### Dobutamine stress measurements

Dobutamine was infused via a peripheral venous cannula, as necessary (typical dosing between 15-40 μg/kg/min) to achieve a target heart rate of 65 % age maximum (calculated as 220-age). ^31^P-MRS and long axis cine images were repeated at maximum stress to elicit energetic and contractile response to stress, as well as exclude any inducible regional wall motion abnormalities.

### Plasma metabolomic and proteomic analyses

Metabolite profiling was performed using standard liquid chromatography-tandem mass spectrometry (LC-MS). Water-soluble metabolites were profiled in the positive ionization mode using an LC-MS system composed of a Nexera X2 U-HPLC (Shimadzu Corp; Marlborough, MA) coupled to a Q Exactive mass spectrometer (Thermo Fisher Scientific; Waltham, MA) and equipped with a hydrophilic interaction liquid chromatography column (Atlantis HILIC; Waters; Milford, MA), as previously described.^29^ Proteomic profiling was performed using the Olink 96 Cardiovascular panel (Olink Proteomics AB, Uppsala, Sweden) according to the manufacturer’s instructions using separate aliquots. The Proximity Extension Assay technology used for the Olink protocol has been previously described.^30^ Detailed metabolite and protein profiling methods are available in the expanded methods in the *Data Supplement*.

### Statistical considerations

Detecting a treatment difference of 0.2 with a beta of 0.8 and an alpha of 0.05 and taking previous studies with metabolic modulators in our centre^31^ in consideration, the current study was designed and powered to assess cardiac energetics (PCr/ATP) in a minimum of 20 patients at 200mg Ninerafxastat BID and data from non-invasive assessment of cardiac metabolism using hyperpolarized [1-^13^C]pyruvate MRS in a minimum of 8 patients. Paired t-tests were performed to assess for changes in plasma levels of proteins and metabolites among individual patients who received ninerafaxstat. We used the Benjamini-Hochberg procedure to correct for multiple comparisons and a false discovery rate (FDR) below 5% was considered statistically significant.

Statistical analyses were performed using commercial software (SPSS 24, Chicago and GraphPad Prism 9, San Diego). All data is presented as median (IQR) or mean (SD) unless otherwise stated. Determination of statistical significance was assessed by Wilcoxon signed-rank test. Pearson’s correlation and linear regression were used where indicated. To compare the coefficient of regression between before and after the trial, dummy variable regression analysis was performed. Values of *p*<0.05 were considered statistically significant.

## Results

### Patient recruitment and disposition

Patient recruitment took place between May and September 2021. All follow-up visits were completed by December 2021. A total of 28 patients were screened for eligibility, 22 were enrolled. One participant withdrew after 2 days of dosing; another participant did not complete the MR examination at their end of trial (EOT) visit due to claustrophobia.

### Baseline population characteristics

Both cohorts (4- and 8-weeks) were well balanced and selected characteristics of the two cohorts are presented in *Table 1*. The mean age for participants was slightly lower in the 8-week cohort (64 ± standard deviation (SD)8.5 years) compared to the 4-week cohort (72 ± SD 0.5 years) with mostly male participants in both cohorts (75 % in the 4-week vs. 56 % in the 8-week cohort). Median glycated haemoglobin A1c (HbA1c) was elevated in both cohorts (7.6 % [IQR 7.5, 7.8] vs. 6.9 % [IQR 6.6, 8.1]). All except one patient in the 4-week cohort were on stable doses of oral antidiabetic medication, with metformin being the most frequently prescribed agent. Most patients enrolled were on a statin (81 %) and renin-angiotensin-aldosterone system (RAAS) blockade (57 %).

**Table 1:** Patient characteristics at baseline for IMPROVE-DiCE.

|  | 4-Week Cohort | 8-Week Cohort |
| --- | --- | --- |
|  | n=4 | n=16 |
| **Anthropometrics, mean (SD)** |  |  |
| Age (years) | 72 (0.5) | 64 (8.5) |
| Male (% of total) | 3 (75) | 9 (56) |
| Weight (kg) | 100 (13.4) | 95 (12) |
| Body mass index (kg/m<sup>2</sup>) | 33.2 (3.8) | 33.1 (3.8) |
| Systolic blood pressure (mmHg) | 142 (9) | 140 (13) |
| Diastolic blood pressure (mmHg) | 74 (5.4) | 75 (8) |
| Resting heart rate (bpm) | 77 (11) | 70 (10) |
| Heart rate during stress (bpm) | 108 (10) | 110 (6) |
| **Metabolic status, median (IQR)** |  |  |
| HbA1c (%) | 7.6 (7.5, 7.8) | 6.9 (6.6, 8.1) |
| Total cholesterol (mmol/l) | 4.2 (3.8, 4.6) | 4.3 (3.4, 5.4) |
| LDL cholesterol (mmol/l) | 2.6 (2.0, 2.8) | 2.3 (1.5, 3.2) |
| HDL cholesterol (mmol/l) | 1.3 (0.9, 1.4) | 1.2 (1.0, 1.3) |
| Triglycerides (mmol/l) | 1.1 (1.0, 1.5) | 1.6 (1.2, 2.3) |
| Fasting glucose (mmol/l) | 8.9 (8.2, 9.6) | 8.1 (5.8, 9.1) |
| Fasting insulin (pmol/l) | 93 (64, 180) | 85 (51, 100) |
| HOMA-IR | 6.0 (4.3,11.7) | 3.9 (1.8, 6.6) |
| Free Fatty Acids (mmol/l) | 0.9 (0.9, 1.0) | 0.8 (0.7, 1.0) |
| **Cardiac Biomarkers, median (IQR)** |  |  |
|  | n=4 | n=16 |
| NT-pro BNP (ng/ml) | 84 (43, 177) | 82 (61, 147) |
| hs-cTn (ng/l) | 4 (2, 7) | 2 (2, 4) |
| **Concomitant Medications, n (%)** |  |  |
| Oral Hypoglycaemic | 3 (95) | 16 (100) |
| ACE-I / ARB | 2 (50) | 10 (63) |
| Ca<sup>2+</sup>-Channel Blocker | 1 (25) | 8 (50) |
| Statin | 2 (50) | 15 (71) |
| Loop diuretic | 1 (25) | 4 (25) |

ACE-I=angiotensin converting enzyme inhibitor; ARB=angiotensin II receptor blocker; bpm=beats per minute;
Ca<sup>2+</sup>=calcium; HDL=high density lipoprotein; HOMA-IR=homeostatic model assessment for insulin resistance; hs-
cTn=high-sensitive cardiac troponin; LDL=low density lipoprotein

### Myocardial metabolism at baseline

At baseline, we observed impaired resting myocardial energetics (median PCr/ATP 1.6 [IQR 1.4, 2.1]) and myocardial steatosis (MTG 2.2 % [IQR 1.5, 3.2]) typical for patients with cardio-metabolic syndromes.

Results from hyperpolarized [1-^13^C]pyruvate MRS were similar to those previously reported.^18^ The [^13^C]bicarbonate to [1-^13^C]pyruvate ratio, shown previously to linearly correlate with *ex-vivo* measurements of PDH-flux, was lower than the literature normal values, and the [1-^13^C]lactate/pyruvate ratio, reflecting exchange through LDH, was increased. As a marker of balance between glycolytic and oxidative carbohydrate metabolism, the ratio of [^13^C]bicarbonate and [1-^13^C]lactate signals showed a significant reduction in relative carbohydrate oxidation in our patients.

Myocardial [^13^C]bicarbonate/pyruvate was inversely correlated with body weight (r=-0.7; *p*=0.04) and insulin resistance (HOMA-IR, r=-0.63; *p*=0.07). Resting myocardial PCr/ATP was inversely correlated with the [^13^C]bicarbonate/lactate ratio, (r=0.73; *p*=0.03) suggesting that higher levels of glucose oxidation are associated with improved energetics. Consistent with their prime role in determining the rate of myocardial FA-uptake, levels of circulating FFA were significantly correlated with MTG content (r=0.66; *p*=0.003), and with diastolic dysfunction (peak circumferential diastolic strain rate r=-0.50; *p*=0.04).

For the following analyses, all data was included and the 4- and 8 weeks results combined, as treatment effects were evident irrespective of duration of treatment.

### Anthropometrics, biomarkers, proteo- and metabolomics

Following treatment, there was a significant reduction in overall body weight (by 0.8 kg [IQR - 2.7, 0.5], *p*<0.05), total serum cholesterol (from 4.3 mmol/L [IQR 3.4, 5.2] to 4.1 mmol/L [IQR 3.5, 4.8], *p*=0.05) and LDL-cholesterol (2.5 mmol/L [IQR 1.6, 3.0] to 2.1 mmol/L [IQR1.7, 2.7], *p*=0.01). Fasting glucose, FFA, HbA1c, insulin and triglycerides were unchanged (see *Table 2)*.

**Table 2:** Effects of ninerafaxstat on selected anthropometrics for the 4- and 8-week treatment groups combined.

|  | Pre-Treatment | Post-Treatment | *P*-value |
| --- | --- | --- | --- |
|  | n=21 | n=21 |  |
| **Anthropometrics, mean (SD)** |  |  |  |
| Age (years) | 66 (8) | 66 (8) | ns |
| Male (% of total) | 12 (57) | 12 (57) | ns |
| Weight (kg) | 96 (12) | 95 (12) | **0.04** |
| Body mass index (kg/m<sup>2</sup>) | 33.6 (3.6) | 33.3 (3.8) | 0.06 |
| Body surface area (m<sup>2</sup>) | 2.05 (0.21) | 2.05 (0.19) | 0.36 |
| Systolic blood pressure (mmHg) | 142 (15) | 140 (13) | 0.42 |

|  | Pre-Treatment<br>n=21 | Post-Treatment<br>n=21 | *P*-value |
| --- | --- | --- | --- |
| Diastolic blood pressure (mmHg) | 73 (8) | 75 (8) | 0.58 |
| Resting heart rate (bpm) | 75 (11) | 70 (10) | **0.009** |
| Heart rate during stress (bpm) | 112 (10) | 110 (6) | 0.31 |
| **Metabolic status, median (IQR)** |  |  |  |
| HbA1c (%) | 7.1 (6.6, 7.9) | 7.2 (6.7, 8.0) | 0.41 |
| Total cholesterol (mmol/l) | 4.3 (3.4, 5.2) | 4.1 (3.5, 4.8) | 0.05 |
| LDL cholesterol (mmol/l) | 2.5 (1.6, 3.0) | 2.3 (1.7, 2.7) | **0.01** |
| HDL cholesterol (mmol/l) | 1.2 (1.0, 1.3) | 1.2 (0.9, 1.3) | 0.99 |
| Triglycerides (mmol/l) | 1.6 (1.1, 2.2) | 1.3 (1.0, 2.4) | 0.69 |
| Fasting glucose (mmol/l) | 8.5 (6.1, 9.1) | 7.2 (5.9, 9.0) | 0.65 |
| Fasting insulin (pmol/l) | 85 (57, 100) | 83 (58, 107) | 0.66 |
| HOMA-IR | 4.1 (2.3, 6.6) | 4.2 (2.9, 6.0) | 0.94 |
| Free Fatty Acids (mmol/l) | 0.9 (0.8, 1.0) | 0.9 (0.5, 1.0) | 0.77 |
| **Cardiac Biomarkers, median (IQR)** |  |  |  |
| NT-pro BNP (ng/ml) | 82 (59, 147) | 60 (40, 138) | 0.70 |
| hs-cTn (ng/l) | 2 (2, 5) | 2 (2, 4) | 0.27 |

ACE-I=angiotensin converting enzyme inhibitor; ARB=angiotensin II receptor blocker; bpm=beats per minute; Ca<sup>2+</sup>=calcium; HDL=high density lipoprotein; HOMA-IR=homeostatic model assessment for insulin resistance; hs- cTn=high-sensitive cardiac troponin; LDL=low density lipoprotein

Plasma metabolomics showed a significant increase of 1-methylnicotinamide (1-MNA, FDR-adjusted p-value<0.05), N1-methyl-2-pyridone-5-carboxamide and niacinamide which are downstream metabolites of niacin, an active moiety of Ninerafaxstat (see *Figure 2 A*).

**Figure 2:**
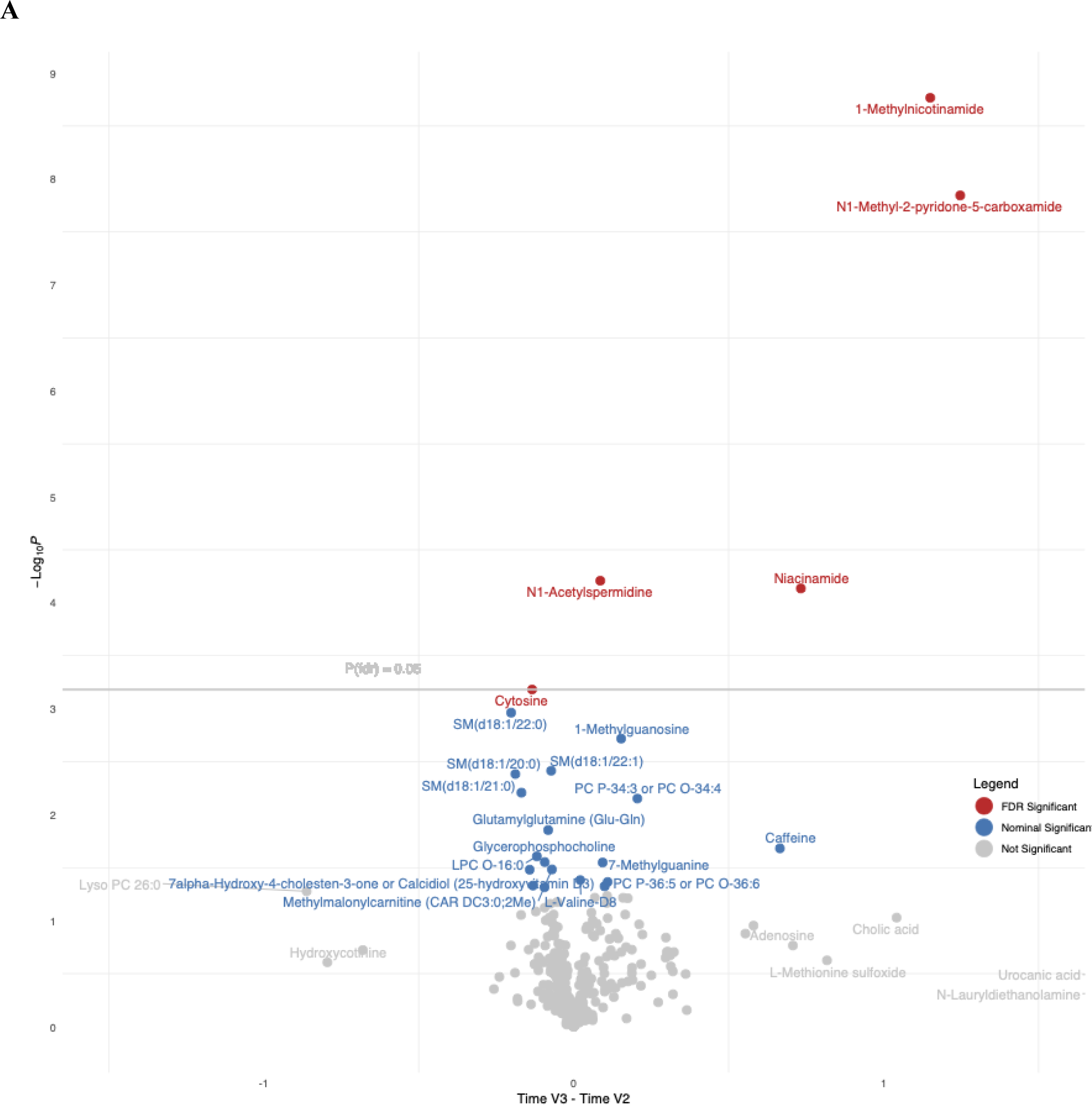
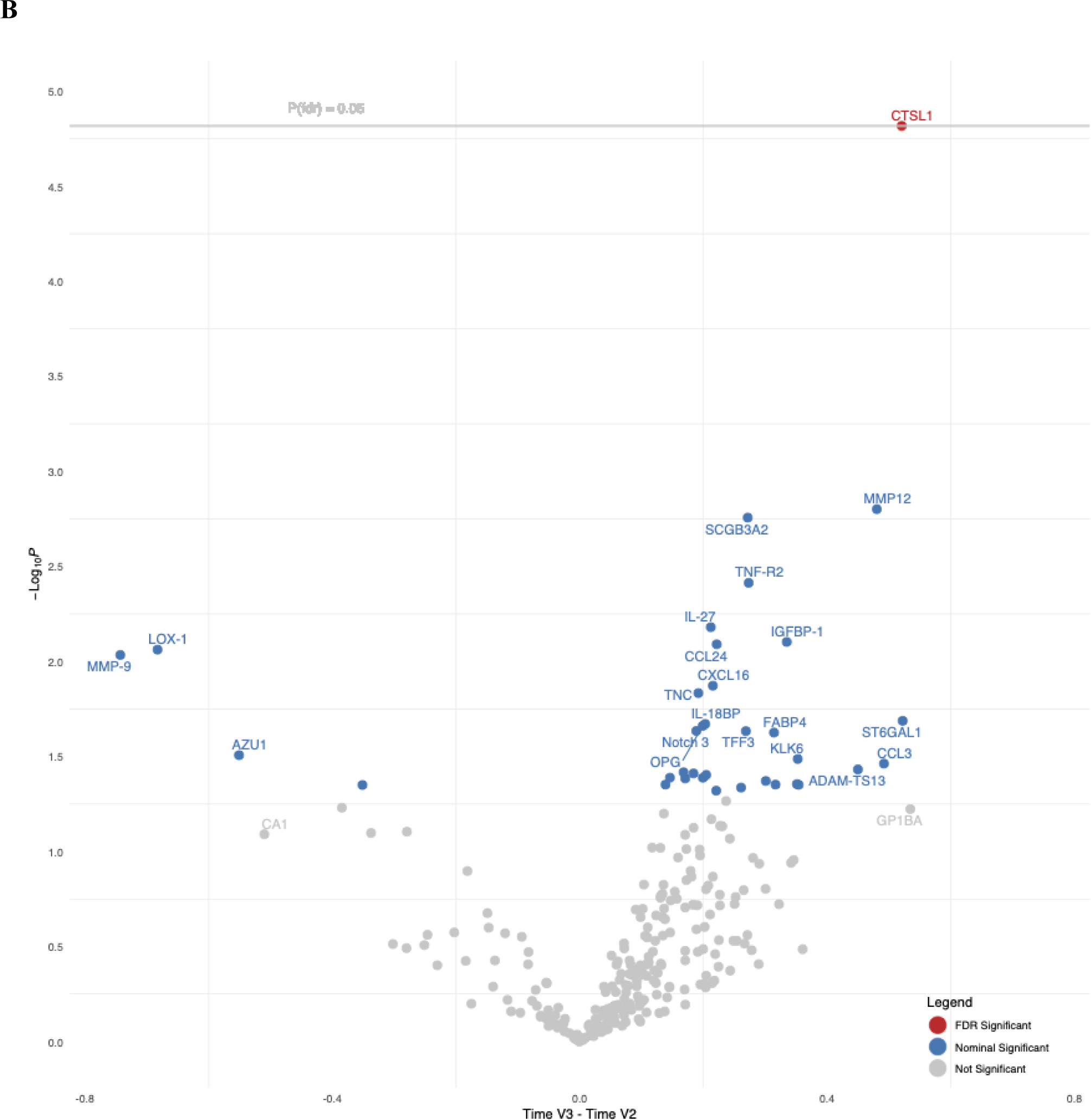
Effects of ninerafaxstat on the plasma metabolome and proteome. Volcano plots of results from plasma metabolomic (A) and proteomic (B) analyses post (V3) vs. pre (V2) treatment with ninerafaxstat for 4- or 8-weeks. ninerafaxstat treatment significantly increased metabolites (A) of the NAD^+^-degradation pathway, confirming its mechanism of action. Proteomic analysis elicited increased circulation of CTSL1 with a modest effect size of other proteins. CTSL1=cathepsin L1; FDR=false discovery rate, NAD^+^=Nicotinamide adenine dinucleotide

Our proteomic analyses revealed a significant upregulation of cathepsin L1 (CTSL1; FDR-adjusted p-value<0.05) while a set of proteins relating to cardiac remodelling (lectin-like oxidized low-density lipoprotein receptor 1), inflammation (matrix metalloproteinase 9) and diastolic dysfunction (fatty acid binding protein 4) showed a trend towards downregulation after treatment (see *Figure 2 B*).

### Cardiac energetics and myocardial steatosis

Ninerafaxstat treatment led to significantly improved myocardial energetics at rest, with a 32 % increase in PCr/ATP (0.13; 95 % confidence interval (CI) 0.1 to 0.5, p<0.01; *Table 4* and *Figures 3 A and 5 C*).

**Figure 3:**
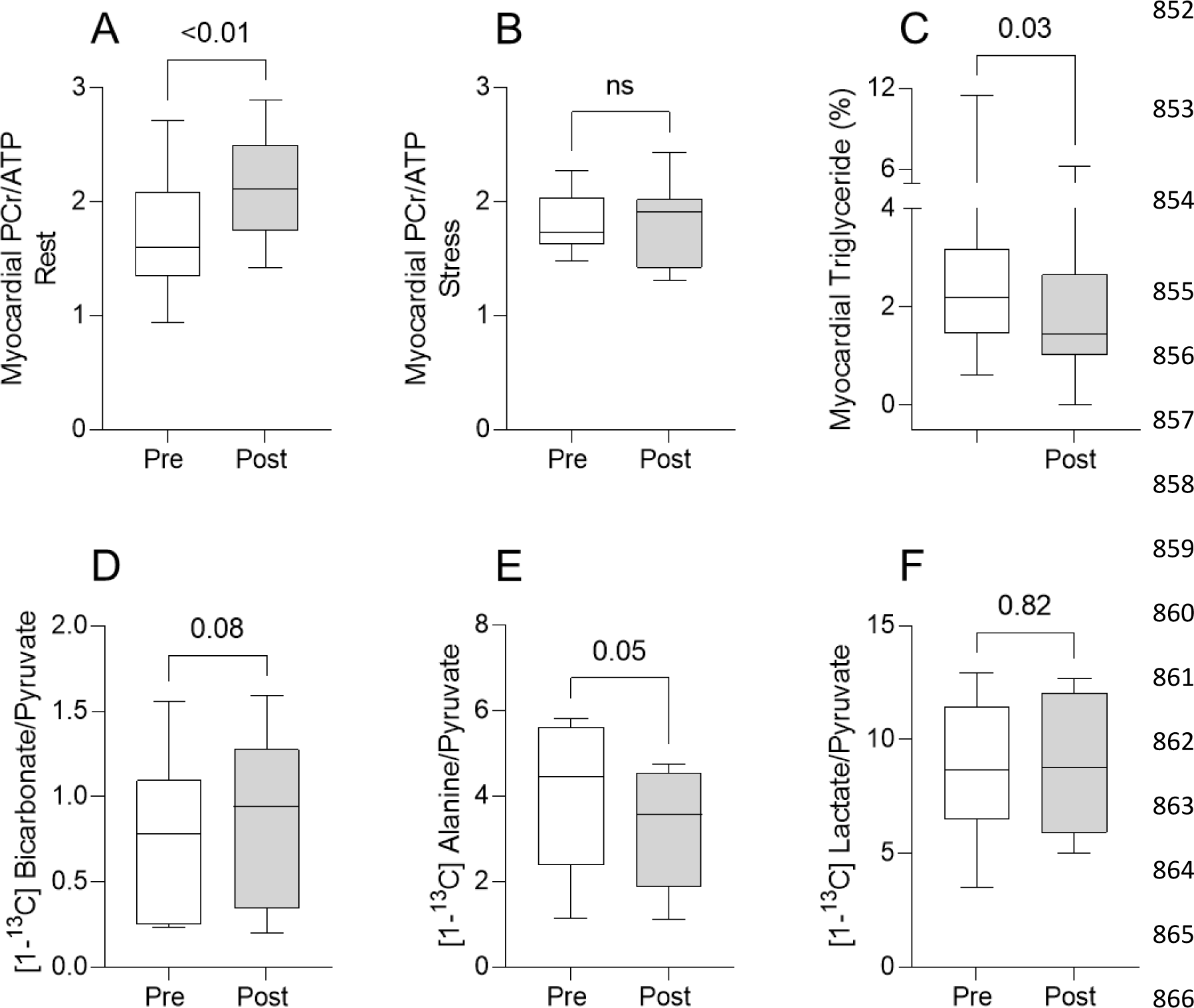
Effects of ninerafaxstat on cardiac energetics and metabolism. Effects of ninerafaxstat on cardiac metabolism (A) resting myocardial PCr/ATP, (B) myocardial PCr/ATP during dobutamine stress, using ^31^P magnetic resonance spectroscopy (C) myocardial triglyceride content (proton density fat fraction, %) recorded with ^1^H magnetic resonance spectroscopy, and (D-F) pyruvate metabolism to bicarbonate, alanine and lactate, respectively, using hyperpolarized [1-^13^C]pyruvate magnetic resonance spectroscopy. Panel (D) denotes one-tailed p-value.

**Figure 4:**
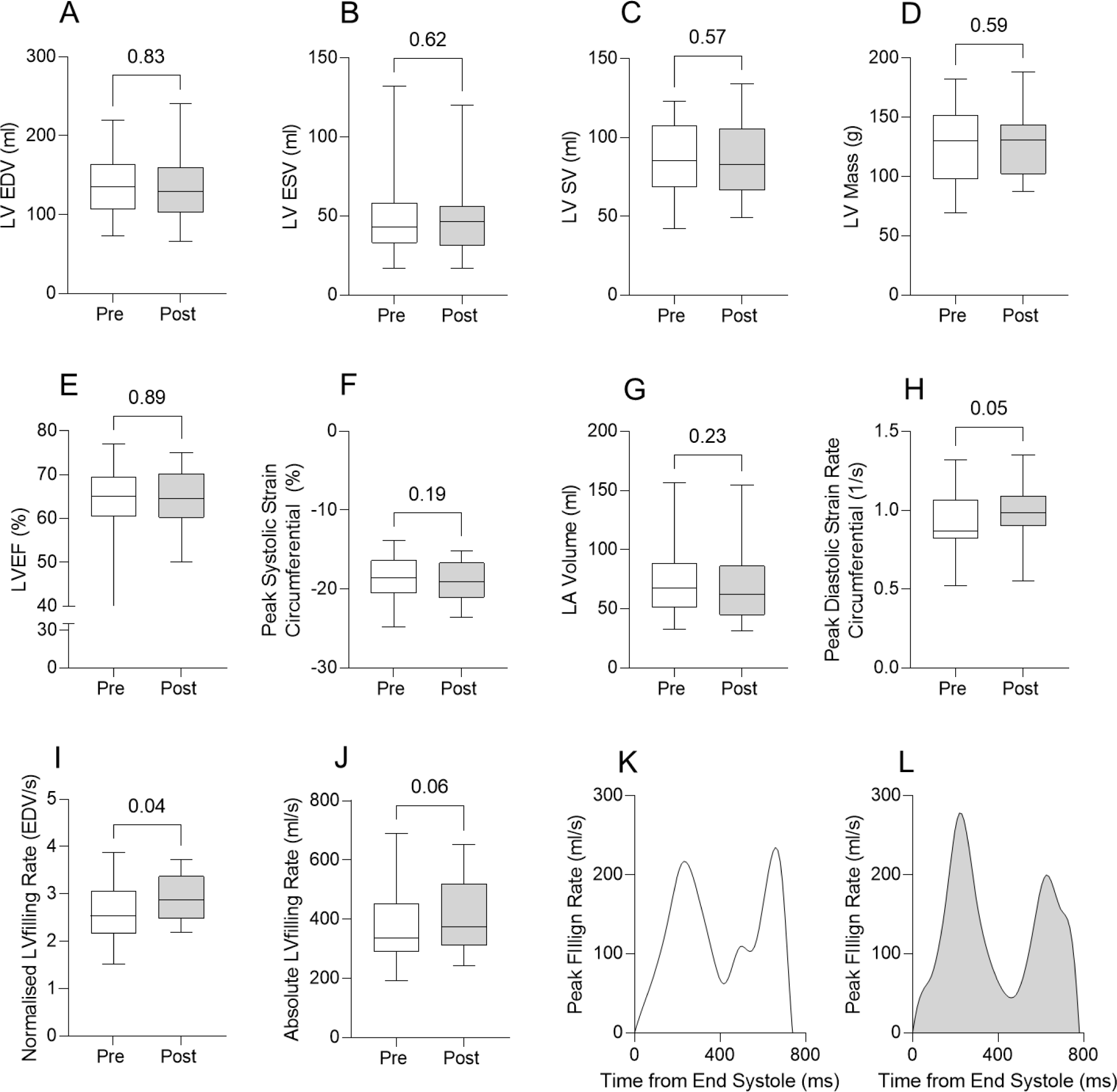
Effects of ninerafaxstat on cardiac structure and function. Effects of ninerafaxstat on left ventricular (LV) structure and systolic function at rest. (A) LV end-diastolic volume (ml), (B) end-systolic volume (ml) (C) stroke volume (ml), (D) LV mass (g), (E) Left Ventricular Ejection Fraction (%), (F) peak systolic circumferential strain (%), (G) LA volume (ml), (H) peak diastolic circumferential strain rate (1/s), (I) normalises and (J) absolute peak LV filling rates, and visual representation of group mean (K) normalised and (L) absolute peak LV filling rates across the cardiac cycle.

**Figure 5:**
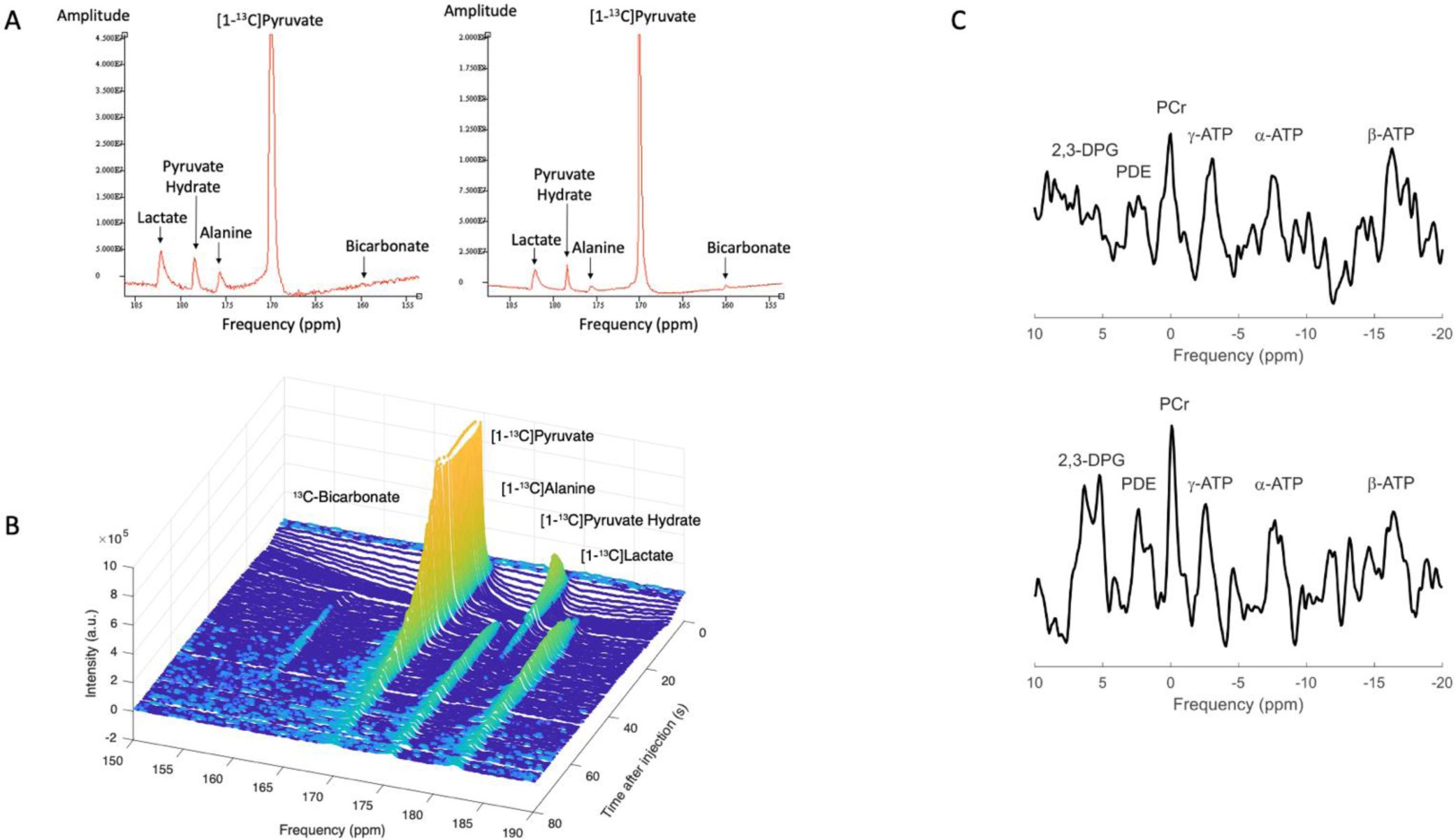
Visualisation of pre- and post-treatment metabolic readouts using [1-^13^C]pyruvate hyperpolarized- and ^31^P-MRS. Representative pre- and post-treatment spectrum acquired with [1-^13^C]pyruvate hyperpolarized MRS (A) and waterfall visualisation displaying all acquired spectra following injection of the hyperpolarized [1-^13^C]pyruvate over time (B). Representative pre- and post-treatment ^31^P-MRS spectra showing an increase in myocardial energy reserve (PCr/ATP) following treatment with ninerafaxstat.

Paired dobutamine stress energetic measurements showed no significant reduction in myocardial PCr/ATP compared to resting values pre-treatment (rest 1.6 [IQR 1.4, 2.1] vs. stress 1.7 [IQR 1.6, 2.0]). Notably, the significant increase in resting myocardial energetics following treatment with ninerafaxstat was constant during stress (PCr/ATP at rest 2.1 [IQR 1.7, 2.5] vs. stress 1.9 [IQR 1.4, 2.0]), with the post-treatment stress myocardial PCr/ATP exceeding pre-treatment resting energetic values. There was no significant difference in stress PCr/ATP between the two timepoints (0.0; 95 % CI −0.2 to 0.2; *p*=0.94; *Table 4*, *Figure 3 B*). As a measure of internal validity, the achieved peak-HR did not differ pre-vs. post-treatment (mean ± SD: pre-treatment 112 ± 5.8 /min vs. post-treatment 110 ± 6.1 /min, *p*=0.31).

MTG content, a measure of myocardial steatosis, was significantly reduced by 34% following treatment with ninerafaxstat (−0.6; 95 % CI −1.3 to −0.2, *p*=0.03, *Figure 3 C*). Whilst fasting FFA-levels remained unchanged following treatment, MTG content remained significantly correlated with FFA-levels (r=0.53, *p*= 0.03), with the coefficient of regression between FFA- and MTG-content being greater post-treatment (+ 7.5 % vs + 1.6% increase per mmol/l increase in FA-level, p<0.01).

### Hyperpolarized ^13^C MRS

As a measure of enzymatic PDH-flux, the [1-^13^C]bicarbonate/pyruvate ratio was, in absolute terms, increased in 7 out of 9 participants, and overall by 20 % (+0.17; 95 % CI −0.03 to 0.5; p=0.08, *Table 4* and *Figure 2D*) not reaching nominal significance. Likewise, the [1-^13^C]alanine/pyruvate ratio was reduced by 20 % following treatment (p=0.05, *Figure 2E*). The [1-^13^C]lactate/pyruvate ratio (*Figure 2F*) and the ratio of [^13^C]bicarbonate/lactate signals remained unchanged with treatment (*p*=0.82, and *p*=0.36 respectively; *Table 4*).

### Cardiac structure and systolic function

There were no changes in LV end-diastolic volume (EDV) or mass (*Table 3, Figure 4*). No change in LV-systolic function was observed during the trial (*Table 3, Figure 4*). RV-size and systolic function likewise did not change post-treatment (*Table 3*).

**Table 3.**
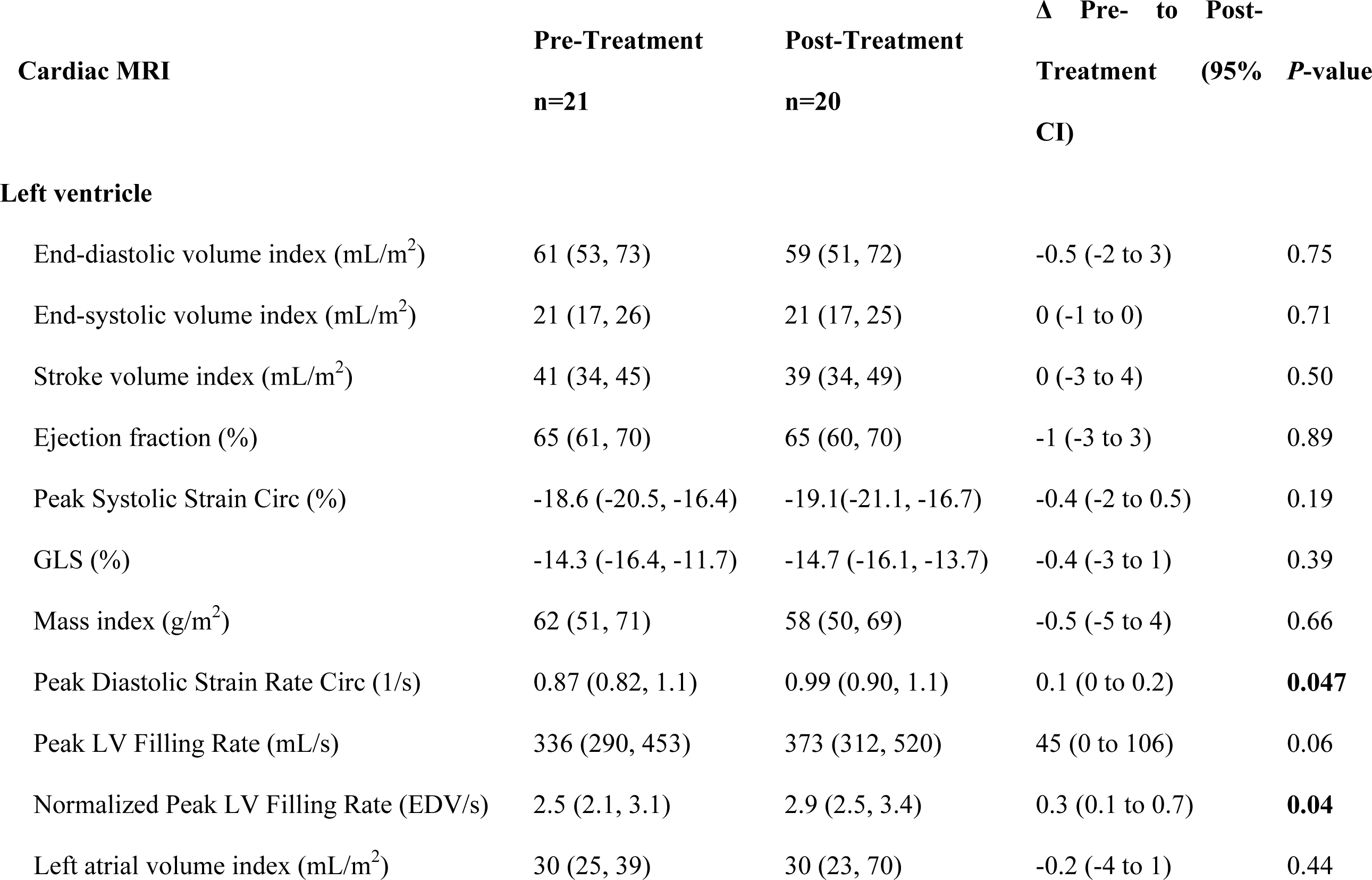
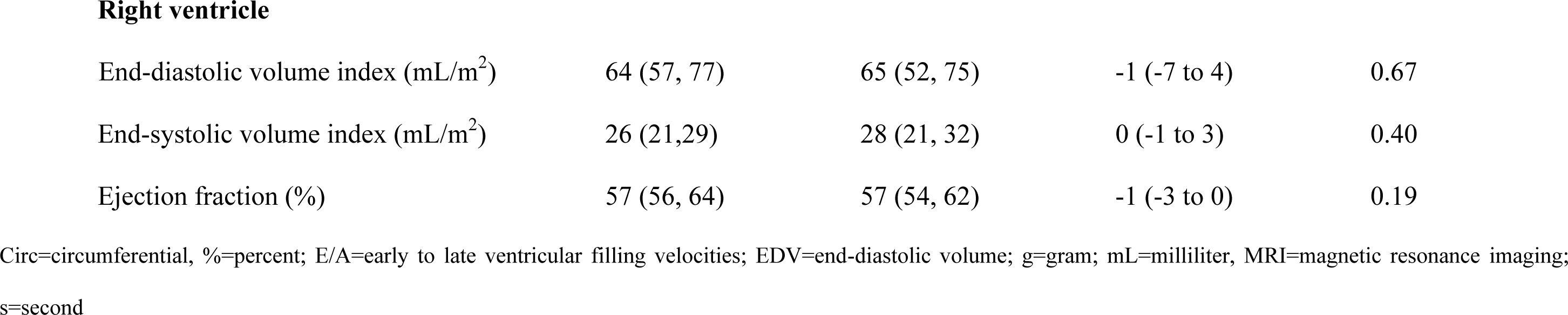
Effects of ninerafaxstat treatment on cardiac structure and function.

| Cardiac MRI | Pre-Treatment<br>n=21 | Post-Treatment<br>n=20 | $\Delta$ Pre- to Post-<br>Treatment (95% CI) | P-value |
| --- | --- | --- | --- | --- |
| **Left ventricle** |  |  |  |  |
| End-diastolic volume index (mL/m<sup>2</sup>) | 61 (53, 73) | 59 (51, 72) | -0.5 (-2 to 3) | 0.75 |
| End-systolic volume index (mL/m<sup>2</sup>) | 21 (17, 26) | 21 (17, 25) | 0 (-1 to 0) | 0.71 |
| Stroke volume index (mL/m<sup>2</sup>) | 41 (34, 45) | 39 (34, 49) | 0 (-3 to 4) | 0.50 |
| Ejection fraction (%) | 65 (61, 70) | 65 (60, 70) | -1 (-3 to 3) | 0.89 |
| Peak Systolic Strain Circ (%) | -18.6 (-20.5, -16.4) | -19.1(-21.1, -16.7) | -0.4 (-2 to 0.5) | 0.19 |
| GLS (%) | -14.3 (-16.4, -11.7) | -14.7 (-16.1, -13.7) | -0.4 (-3 to 1) | 0.39 |
| Mass index (g/m<sup>2</sup>) | 62 (51, 71) | 58 (50, 69) | -0.5 (-5 to 4) | 0.66 |
| Peak Diastolic Strain Rate Circ (1/s) | 0.87 (0.82, 1.1) | 0.99 (0.90, 1.1) | 0.1 (0 to 0.2) | **0.047** |
| Peak LV Filling Rate (mL/s) | 336 (290, 453) | 373 (312, 520) | 45 (0 to 106) | 0.06 |
| Normalized Peak LV Filling Rate (EDV/s) | 2.5 (2.1, 3.1) | 2.9 (2.5, 3.4) | 0.3 (0.1 to 0.7) | **0.04** |
| Left atrial volume index (mL/m<sup>2</sup>) | 30 (25, 39) | 30 (23, 70) | -0.2 (-4 to 1) | 0.44 |
| **Right ventricle** |  |  |  |  |
| End-diastolic volume index (mL/m<sup>2</sup>) | 64 (57, 77) | 65 (52, 75) | -1 (-7 to 4) | 0.67 |
| End-systolic volume index (mL/m<sup>2</sup>) | 26 (21,29) | 28 (21, 32) | 0 (-1 to 3) | 0.40 |
| Ejection fraction (%) | 57 (56, 64) | 57 (54, 62) | -1 (-3 to 0) | 0.19 |

Circ=circumferential, %=percent; E/A=early to late ventricular filling velocities; EDV=end-diastolic volume; g=gram; mL=milliliter, MRI=magnetic resonance imaging;
s=second

**Table 4.**
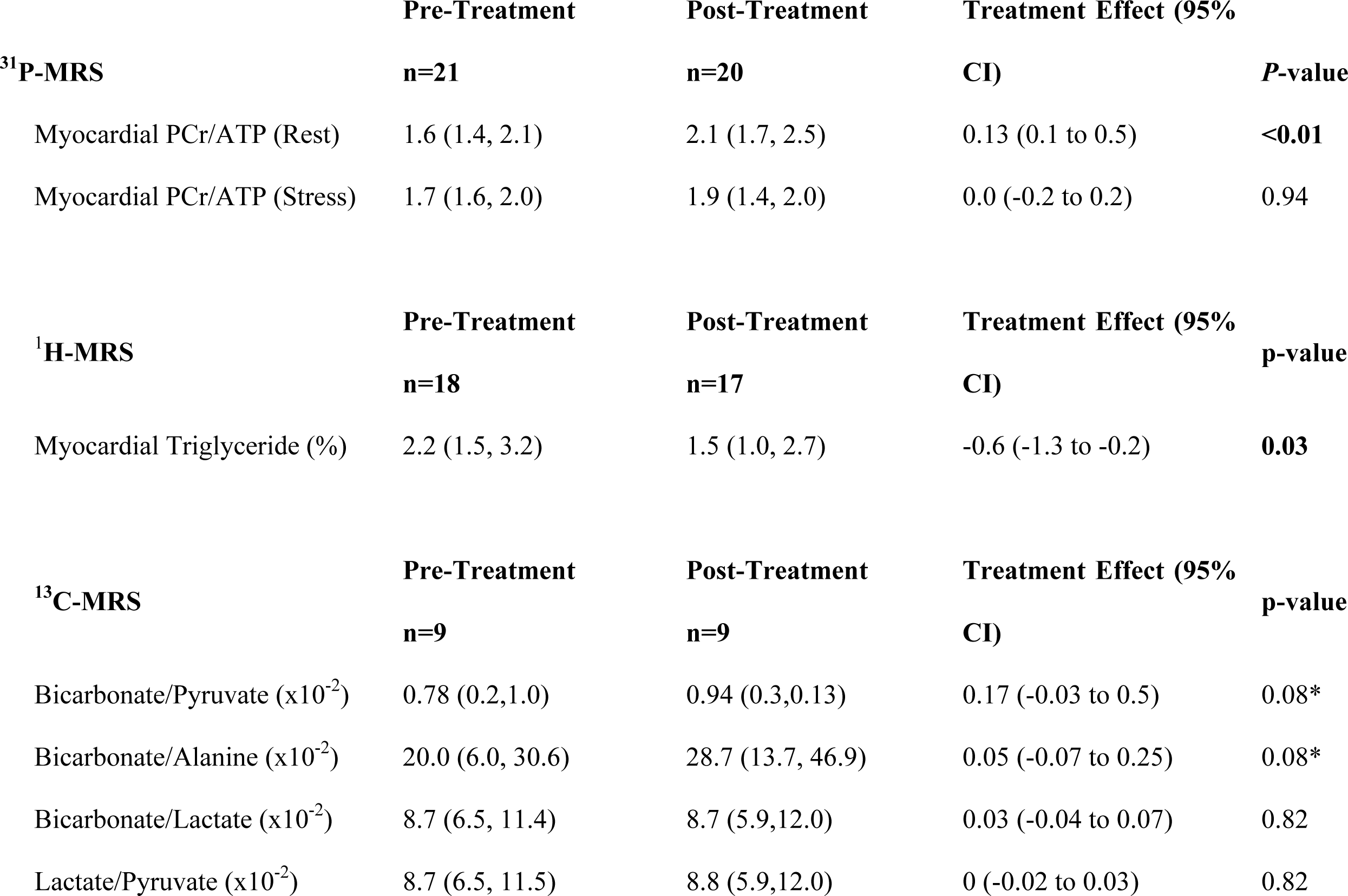
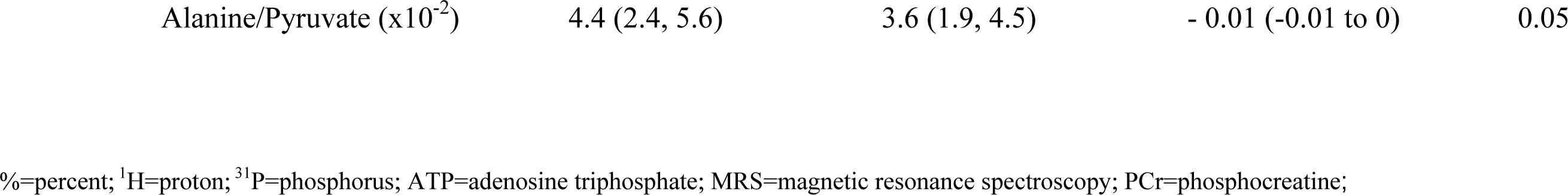
Effects of ninerafaxstat on cardiac metabolism using ^31^P, ^1^H and hyperpolarized [1-^13^C] MRS, *denotes one-tailed P value.

### LV diastolic function

Following treatment, there was a 15 % increase in peak diastolic strain rate (+0.1; 95 % CI 0 to 106; *p*<0.05, *Table 3*, *Figure 3H*) supported by a significantly increased peak LV filling rate normalised to EDV (0.3 EDV/s; 95 % CI 0.1 to 0.7; p=0.04). There were no changes in left atrial volume index (−1 mL/m^2^, 95 % CI −3 to 0; *p*=0.44; *Table 3*).

### Safety and tolerability

Ninerafaxstat was safe to use and well tolerated. Of 22 participants, 13 (∼59 %) reported 15 treatment-emergent adverse events (AEs) all of which were described as self-limiting and mild or moderate in severity. 5 AEs (1 mild, 4 moderate) were judged to be related to study drug with one participant developing diarrhoea and dizziness (both moderate in severity) after 2 days of dosing, resulting in elected withdrawal from the study.

There were no treatment-emergent serious AEs (SAEs), dose modifications or suspension of dosing.

## Discussion

The ‘diabese’ (T2D+obesity) heart is characterised by metabolic remodelling including increased FA uptake and high levels of FAO with reduced PDH-flux, ultimately resulting in lipotoxicity.^18^ On a cellular level, the excessive reliance on FAO for ATP-generation results in an energy deficit and cardiac steatosis, LV hypertrophy and LV diastolic dysfunction.^32^

In this mechanistic trial, we evaluated the mechanism of action and impact of the novel metabolic modulator ninerafaxstat, on myocardial energetics, metabolism and function in participants with cardio-metabolic syndromes. Using ^31^P-MRS, we observed that 4-8 weeks treatment are sufficient to improve myocardial energetics (PCr/ATP). Using hyperpolarized [1-^13^C]pyruvate MRS, for the first time in a clinical trial of a novel cardiac IMP, we observed cumulative changes consistent with increased PDH-flux. Furthermore, ninerafaxstat reduced myocardial steatosis and we demonstrate that these modifications of myocardial metabolism and energetics translate into improved diastolic function, showcasing the potential of CMR to investigate novel compounds for treatment of patients with cardio-metabolic syndromes.

### Myocardial energetics

Myocardial high-energy phosphate metabolism plays a crucial role in maintaining normal systolic and diastolic cardiac function and homeostasis. The heart’s energetic status, expressed as PCr/ATP, is reduced across the spectrum of HF, from sub-clinical patients with T2D/obesity at risk for HF ^33^ to clinically overt HFpEF.^34^ Myocardial PCr/ATP has been shown to independently predict both all-cause and cardiovascular (CV) mortality in patients with dilated cardiomyopathy and to correlate with the clinical severity of HF (NYHA functional class), LVEF and LV wall thickness.^35^

Following 4-8 weeks of ninerafaxstat treatment, myocardial PCr/ATP improved significantly, with the median PCr/ATP at the end-of-treatment recovering to the physiologically normal range, consistent with enhanced cardiac energy reserves.^32^ Via the creatine kinase (CK) reaction, an increase in PCr is expected to support myocardial ATP regeneration at a much faster rate than it is consumed (with ATP turnover far exceeding that of ATP synthesis by glycolysis and oxidative phosphorylation), enabling maintenance of high levels of intracellular ATP and low levels of free ADP, in keeping with continuous free energy release from ATP hydrolysis particularly during stress.

Notably, the absolute magnitude of improvement in PCr/ATP reported here is comparable to or greater than that described with approved guideline-directed medical therapies for HF and/or T2D such as β-blockers^36^, sodium-glucose cotransporter-2 (SGLT2) inhibitors^37^, or perhexiline^38^. Our findings support the hypothesis that metabolic modulation with ninerafaxstat has the potential to enhance myocardial metabolism, resulting in beneficial functional changes in patients with cardio-metabolic syndromes.

### Myocardial triglyceride content

T2D, obesity and HFpEF are associated with increased lipid deposition leading to lipotoxicity.^39^ A progressive gradient of myocardial steatosis has been shown across the spectrum of cardio-metabolic syndromes.^40^ Myocardial steatosis in T2D and obesity is independently associated with concentric LV-remodelling, contractile dysfunction^32^ and diastolic dysfunction.^41^

Using ^1^H-MRS, our cohort exhibited marked myocardial steatosis and showed near universal reduction in MTG after 4-8 weeks of ninerafaxstat (see *Table 3*). Whilst the MTG storage pool is dynamic, relatively few pharmacological therapeutic interventions have been demonstrated to impact upon it. Prolonged caloric restriction in the form of a very-low-energy diet (VLED, 450 kcal/day) for 16 weeks in individuals with T2D and obesity reduced MTG (from 0.88 ± 0.12 to 0.64 ± 0.14 %, mean ± standard error (SE)) with a marked fall in BMI (from 35.6 ± SE 1.2 to 27.5 ± SE 1.3 kg/m^2^) and improved diastolic function.^42^ Similarly, our group have shown a fall in MTG content (from 2.1 ± 1 to 1.5 ± 0.6 %) after 8 weeks of a VLED (800 kcal/day) in patients with obesity (BMI 36.8 ± 5.8 kg/m^2^).^43^ Whilst such substantial reductions in body weight have also been shown to improve myocardial energetics^44^ and LV diastolic function^45^, we observed only a modest fall in body weight in participants during the study (median difference pre-vs. post-treatment 0.8 kg, *p*<0.05) which was far smaller than would be expected to drive the magnitude of reduction in MTG (or improvement in energetics) observed.

The mechanisms underlying the simultaneous reduction of myocardial FAO (as an increase in glucose oxidation would support) whilst markedly reducing MTG in the absence of an overt impact on circulating FFA remain to be elucidated. Short-term administration of TMZ has been reported to inhibit both total and endogenous myocardial FAO in obese participants, although without a significant change in lipid content when assessed after 4 weeks treatment.^46^ In the present study, this was demonstrated by a reduction in MTG content without change in circulating FFA levels in combination with a stronger correlation between FFA and MTG content before treatment.

### PDH flux

The inhibitory effect of increased FA availability on glucose oxidation and competition for acetyl CoA for the tricarboxylic acid cycle (the ‘glucose-fatty acid cycle’ or ‘Randle cycle’) have been recognised for over half a century.^47^ Whilst the ability to partition limited glucose for use by glycolytic tissues has been proposed to confer an evolutionary advantage in the face of starvation (which leads to increased circulating FFA), in the context of a chronic caloric surplus and fat-rich nutrient diet, the inability to enhance glucose metabolism to meet myocardial energy demands is likely maladaptive. Consequently, increasing PDH-flux has been proposed as a therapeutic strategy to improve cardiac energetics and function across a range of myocardial disease states characterised by dynamic metabolic inflexibility and impaired glucose oxidation (T2D, obesity, obese HFpEF).^48^

Using hyperpolarized [1-^13^C]pyruvate MRS, we observed a numerical increase in the [^13^C]bicarbonate/ [1-^13^C]pyruvate ratio in 7 of 9 participants treated with ninerafaxstat (by 20 %, one-sided *p*=0.08). Moreover, we detected a strong signal for reduced transamination of pyruvate to alanine, inferred from the change in [1-^13^C]alanine/pyruvate ratio) [by ∼18 %, *p*=0.05], without evidence of impact on exchange through LDH. Together, these results are consistent with increased enzymatic flux through PDH and enhanced myocardial glucose oxidation via inhibition of FAO, restoring the balance of substrate use in the diabetic myocardium.

One of Ninerafaxstat’s active ingredients is Trimetazidine (TMZ), which is a partial FAO-inhibitor that has previously been found to increase total myocardial glucose utilisation (i.e. oxidative and glycolytic), ATP-content and cardiac output^49^ and to reduce myocardial FAO with unchanged overall oxidative metabolism (consistent with increased oxidation of glucose).^50^ Patients with HF displayed increased myocardial energetics, LVEF and improved NYHA class and exercise capacity following TMZ-treatment.^51^ Thus, our results further validate metabolic modulation as noteworthy therapeutic targets which translate to measurable functional cardiac changes using CMR.

### Plasma –‘omics’

An increasing amount of literature has formed a detailed picture of metabolite and protein signatures of patients at risk for or with established HF.^52–54^

Utilising metabolimic and proteomic profiling, our results support Ninerafaxstat’s mechanism of action by showing an upregulation of metabolites involved in the degradation of nicotineamide adenine dinucleotide (NAD^+^) via the amidation pathway. NAD^+^ repletion has recently been shown to improve mitochondrial function and reverse metabolic remodelling in HFpEF, specifically via enhancing PDH-flux.^55^ This is in keeping with our findings of improved cardiac energetics, diastolic function and the observed improvedment of PDH-flux in 7 of 9 patients. Additionally, higher levels of N1-Acetylspermidine, as shown in *Figure 2A*, corresponded with a significant reduction of incident HF in a population at risk.^29^

Despite a modest effect size, our proteomic analyses elicited a CTSL-1-upregulation following treatment. This has recently been identified as possibly protective towards development of HF and was suggested as a possible therapeutic target for treatment of HF.^52^ Additionally, it was shown to attenuate cardiac hypertrophy via activation of lysosomal degradation pathways, which is in keeping with our observation of reduced myocardial steatosis despite inhibition of FAO.^56^

### Diastolic function

Mitochondrial energy and the free energy of ATP-hydrolysis are negatively affected by an over-reliance on FA.^57^ Following treatment, we identified an improved LV-diastolic function, measured by a significantly increased LV peak diastolic strain rate (PDSR) (*p*<0.05) and an increase in normalised LV peak filling rate (*p*<0.05).

### Study Limitations

Using cutting-edge techniques, we provide a detailed assessment of metabolic changes induced by treatment with ninerafaxstat using not only multi-nuclear MRS (^31^P- and ^1^H-) but also hyperpolarized [1-^13^C]pyruvate MRS as well as plasma metabolomics and proteomics. Furthermore, the additional use of MRI, the gold standard imaging modality to assess cardiac function and structure, further increases reproducibility of our findings.

Nevertheless, limitations of our study include its open-label design, recruitment at a single center, the limited sample size and the exclusion of patients on SGLT2i and insulin, as well as those with atrial fibrillation. Female and male participants were not entirely equal in our cohort and ethnic minorities were also underrepresented.

## Conclusions

In patients with cardio-metabolic syndromes, reducing lipotoxicity via partial FAO-inhibition with Ninerafaxstat corrects the energetic impairment of the heart, reduces myocardial steatosis and improves LV-diastolic filling, assessed by CMR. On a broader scale, combining hyperpolarized MRS with metabolic readouts and MRI, showcases the ability of CMR to investigate drug effects *in-vivo*, even with small sample sizes thus, enhancing CV-drug development in early study phases. The metabolic, energetic and functional effects observed, support further investigation of the potential of metabolic modulation to treat or prevent the progression of HF-states associated with diabetes and/or obesity: DbCM and obese HFpEF.

## Novelty and Significance

### What is known?

- Altered myocardial metabolism is a hallmark of virtually all heart diseases. In conditions such as obesity and T2D, disproportionate FAO results in lipotoxicity which ultimately leads to an energetic deficit and cardiac dysfunction.
- Hyperpolarised pyruvate MR was shown to be an effective tool to assess cellular metabolism in real time in-vivo and can be combined with other established metabolic assessments and CMR imaging.
- Given the significant prevalence of cardio-metabolic phenotypes in heart diseases (e.g. HFpEF) and successful metabolic modulation in animal models and other cardiac conditions (e.g. HCM), we aimed to examine effects of promoting glucose usage in lipotoxic conditions in patients with obesity and T2D.

### What new information does this article contribute?

The present mechanistic, phase 2a trial, enrolled 21 participants who were treated with 200mg ninerafaxstat twice daily for 4 (n=5) or 8 weeks (n=16) and assessed by hyperpolarized MR-spectroscopy, CMR and metabolomics pre- and post-treatment. Shifting substrate utilisation towards glucose oxidation, ninerafaxstat treatment improved myocardial energetics, diastolic function and reduced myocardial steatosis. Our results highlight the potential of specific metabolic modulation as novel treatment options for patients with cardio-metabolic HF-phenotypes (such as HFpEF) and corroborate the unique ability of CMR.

## Acknowledgements

The authors gratefully acknowledge the participants in this study for their time and patience, and express their gratitude towards the OCMR research nursing team for their support in conducting this study, especially Mrs. Judith Delossantos and Ms. Harriet Nixon.

## Central Figures

Effects of short-term treatment with ninerafaxstat on cardiac energetics, metabolism and function in patient with T2D and obesity

## List of Nonstandard Abbreviations and Acronyms

^1^H-MRS: proton magnetic resonance spectroscopy
^31^P-MRS: phosphorus magnetic resonance spectroscopy
AE: adverse event
ACE-I: angiotensin converting enzyme inhibitor
ALAT: alanine aminotransferase
ARB: angiotensin II receptor blocker
ATP: adenosine triphosphate
BID: twice daily
BMI: body mass index
CMR: cardiovascular magnetic resonance
DbCM: diabetic cardiomyopathy
EDV: end-diastolic volume
EOT: end of trial
FFA: free fatty acids
FAO: fatty acid oxidation
FT-CMR: feature tracking CMR
GLS: global longitudinal strain
HbA1c: glycated haemoglobin A1c
HLA: horizontal long axis
HF: heart failure
HFpEF: heart failure with preserved ejection fraction
HFrEF: heart failure with reduced ejection fraction
HOMA: homeostasis model assessment
Hs-cTn: high-sensitivity cardiac troponin
IQR: interquartile range
LDH: lactate dehydrogenase
LV: left ventricular
LVEDV: left ventricular end-diastolic volume
LVEF: left ventricular ejection fraction
LVOT: left ventricular outflow tract
MRI: magnetic resonance imaging
MRS: magnetic resonance spectroscopy
MTG: myocardial triglyceride
NAD+: nicotinamide adenine dinucleotide
NT-proBNP: n-terminal pro b-type natriuretic peptide
NYHA: New York Heart Association
PCr/ATP: phosphocreatine to adenosine triphosphate ratio
PDH: pyruvate dehydrogenase
PDSR: peak diastolic strain rate
QUICKI: quantitative insulin sensitivity check
ROS: reactive oxygen species
SAE: serious adverse event
SCMR: Society for Cardiovascular Magnetic Resonance
SGLT2i: sodium glucose co-transporter 2 inhibitor
SV: stroke volume
T2D: type 2 diabetes
TMZ: trimetazidine
UTE: ultra-short echo time
VLA: vertical long axis
VLED: very low energy diet

---

**Figure 6:**
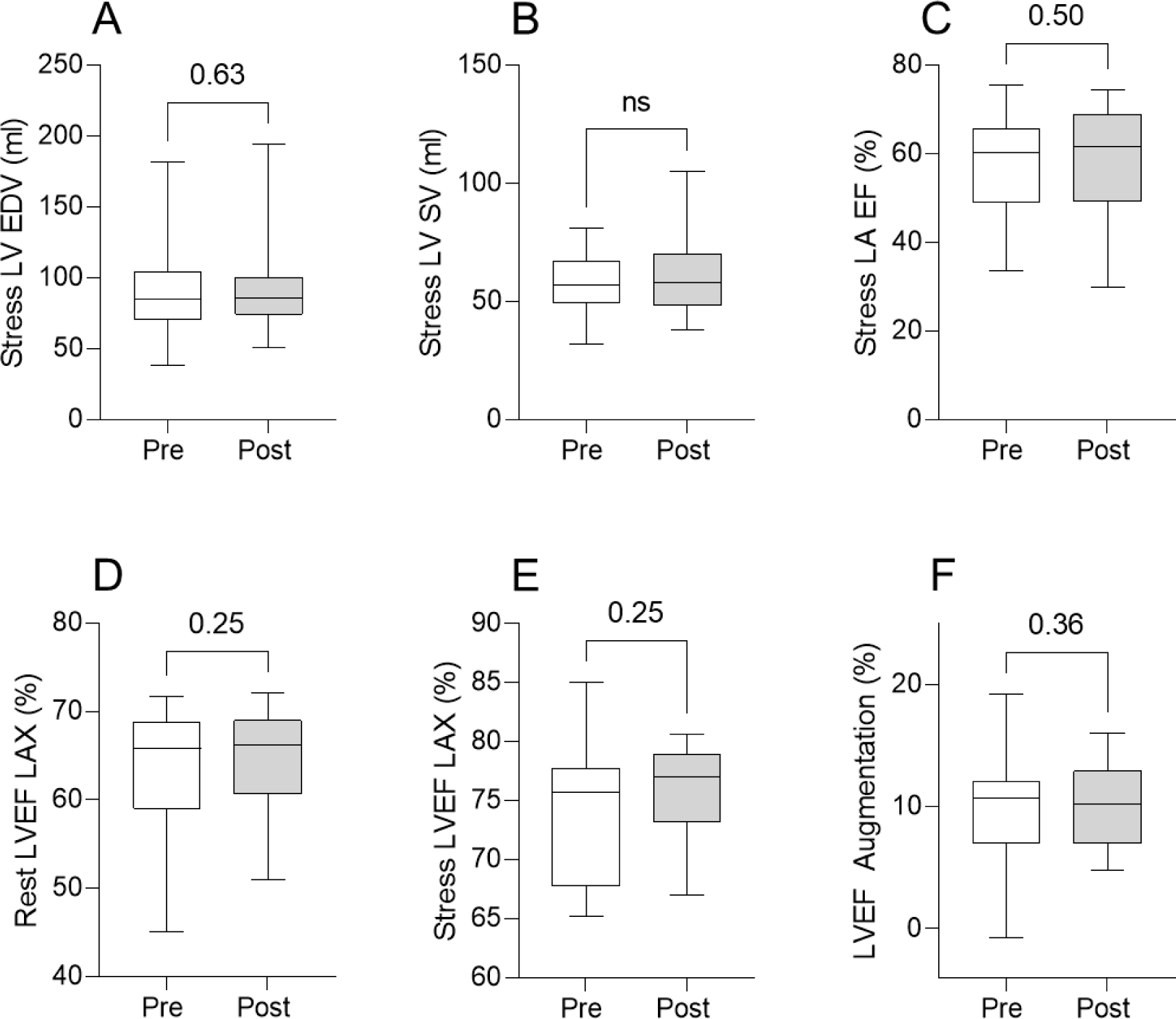
Effects of ninerafaxstat treatment on cardiac function during dobutamine stress. Effects of ninerafaxstat on Cardiac Function during dobutamine stress (A) LV end diastolic volume, (B) LV stroke volume, (C) left atrial ejection fraction (LAEF) spectroscopy, and LV ejection fraction at rest (D), stress (E) and the LVEF augmentation (F). All ventricular measurements for this analysis derived from horizontal and vertical long axis cines.

## Hundertmark et al.: Data Supplement

### 1

**Table S1.**
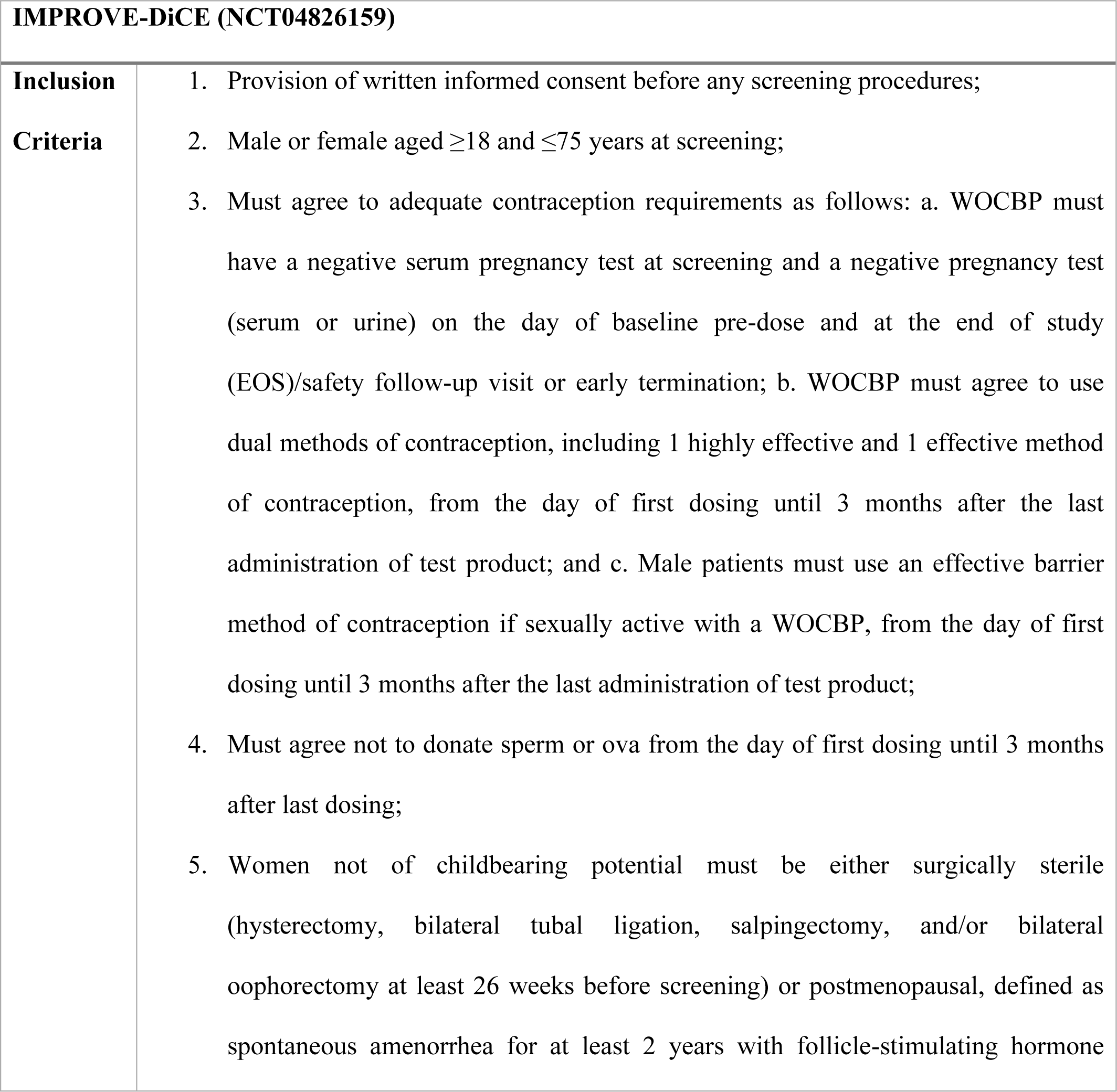
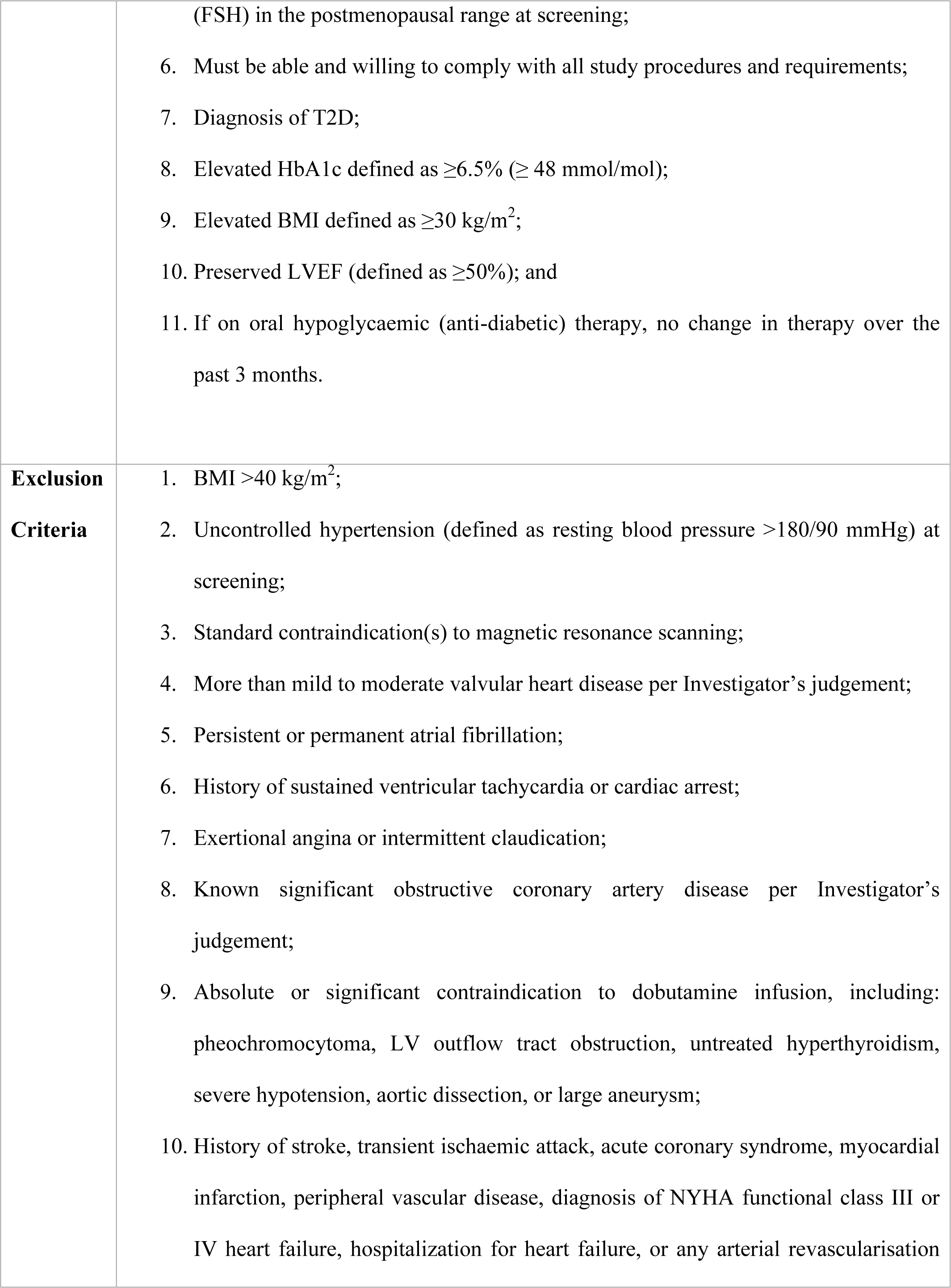
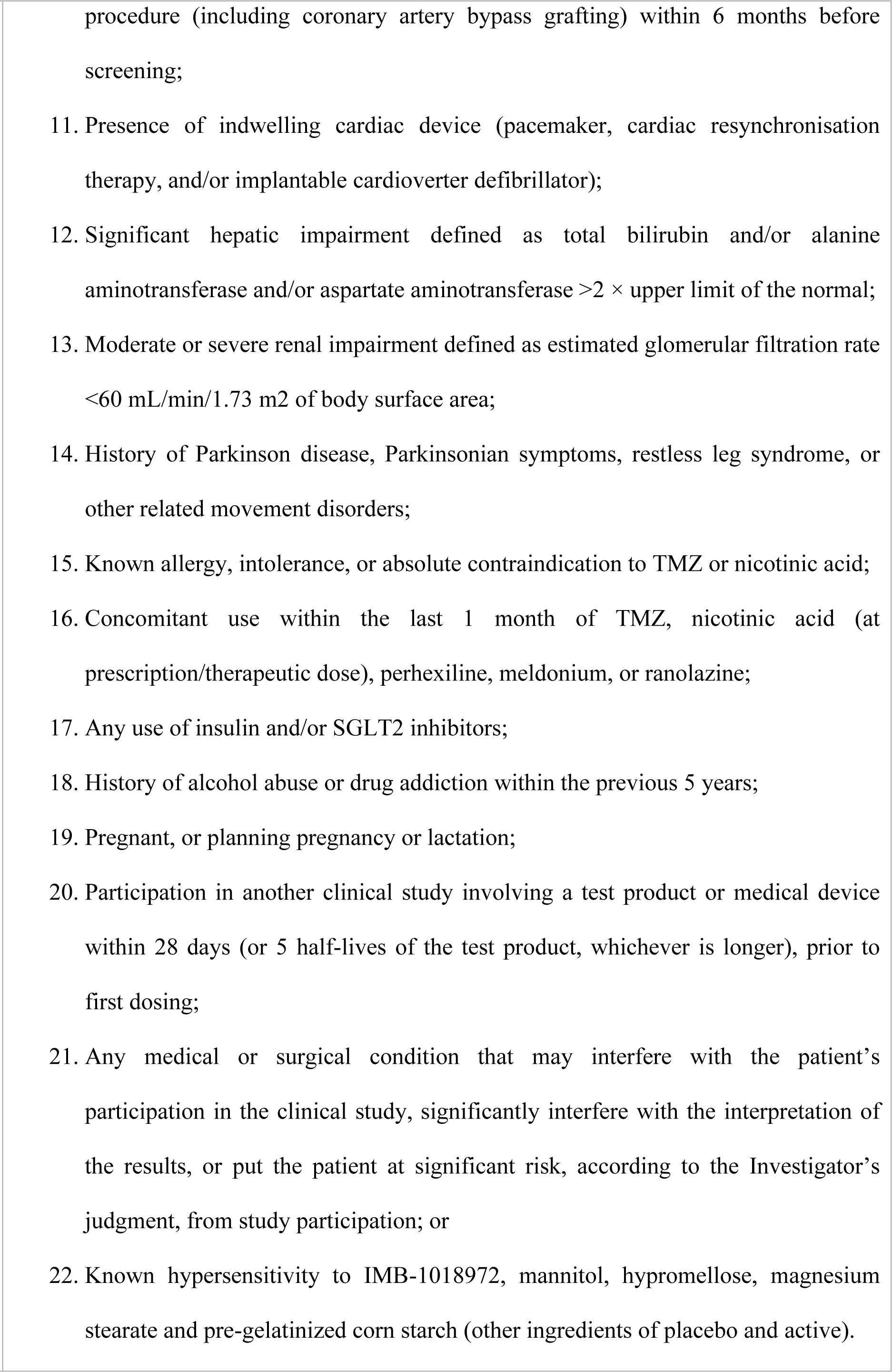
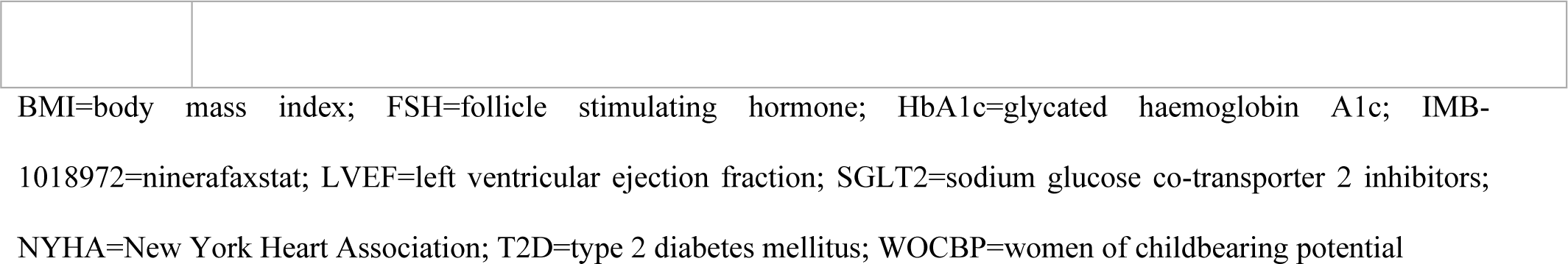
Trial in- and exclusion criteria.

### 2 Hyperpolarized [1-^13^C]pyruvate Production and Injection

Sterile fluid pathways (SFPs) were assembled in a Grade A sterile environment containing 1.47 g [1-^13^C]pyruvic acid (Sigma Aldrich, Gillingham, UK) and 15 mM AH111501 (Syncom, Groningen, Netherlands) as the electron paramagnetic agent (EPA). SFPs were loaded into a General Electric SpinLab system (GE Healthcare, Chicago, USA) which was used for the process of Dynamic Nuclear Polarization. Sufficient polarization levels were achieved after 2-3 hours. Dissolution was undertaken using 38.5 g of sterile water heated to 130°C under pressure, released through the pyruvate containing vial into a receiver vessel containing 17.7 g of trometamol buffer solution (600 mM NaOH, 333 mM Tris base, and 333 mg/L disodium EDTA [as the chelating agent], Royal Free Hospital, London, UK) and a further 19.5 g of sterile water. The EPA was removed by filtration prior to the receiver vessel, with the final product for injection drawn from the receiver vessel into a 50 ml injection syringe (Bayer, Indianola, USA) via a further 0.2 µm sterilization filter (Saint-Gobain, Gaithersburg, USA). Rigorous quality control (QC) of the final filtered sodium [1-^13^C]pyruvate solution was undertaken prior to human injection. This consisted of both online measurements (pyruvate concentration, residual EPA concentration, temperature, polarization, volume) directly from the SpinLab inbuilt QC console, with further ‘offline’ pH measurement (RQflex 10, Merck, Darmstadt, Germany) and visual inspection of the product (for visible particulates and appearance) undertaken manually prior to release. Pathways were only released for human injection if the following criteria were met: pH 6.7-8.4, temperature 25.0-37.0°C, polarization ≥ 15%, [pyruvate] 220-280 mM, [EPA] ≤ 3.0 µM, appearance: clear, colourless solution with no visible particulate matter. Pathways not meeting these release criteria were rejected. Hyperpolarized [1-^13^C]pyruvate solution was administered through an 18G venous cannula sited in the left antecubital fossa, at a dose of 0.4 ml/kg, followed by a 25 ml 0.9% normal saline flush. Injections were performed at a rate of 5 ml per second using a MEDRAD® power injector system (Bayer, Berlin, Germany).

#### Spectral Analysis

Multi-coil data were recombined in MATLAB using the Whitened Singular Value Decomposition algorithm, with coil combination weights calculated for spectra with the highest SNR subsequently applied to the entire dataset. Spectra were background-subtracted prior to quantification with the AMARES algorithm, with appropriate prior knowledge. Total integrated metabolite-to-pyruvate ratios, known to linearly correlate with first-order chemical kinetic rate constants, were calculated from 60 seconds of data taken after the initial appearance of the pyruvate resonance in the spectrum.

### 3 ^31^P-MRS and Data Processing

All scans were performed on a Siemens three Tesla (3T) Tim Trio system (Erlangen, Germany). A Siemens Heart/Liver ^31^P coil was used consisting of a large outer element (26 x 28 cm) which acts as ^1^H transmit-receive and ^31^P transmit, with a smaller loop/butterfly receive pair (12 x 15 cm loop and 23 x 12 cm butterfly) which receives ^31^P signal.

Subjects lay prone with their left ventricle positioned over the centre of the coil at the magnet iso-center. Proton localisers were used to position the subject correctly. Ten free induction decay inversion recovery (IR-FID) curves (1 ms hard inversion) with increasing inversion delay (100 – 3000 ms) are acquired, along with locations of phenylphosphonic acid (PPA) fiducial and codliver-oil phantoms.

Pilot images were taken to position the CSI matrix. Piloting was performed in vertical long axis (VLA), horizontal long axis (HLA) and short axis planes, where a stack of 20 slices was obtained. Fast low-angle shot (FLASH) images were used: slice thickness 10 mm, TR 7 ms, TE 3.37 ms, FOV 400 x 340 mm.

A three-dimensional acquisition-weighted chemical shift imaging (CSI) was used with an acquisition matrix size measuring 16 x 8 x 8 and a field of view of 240 x 240 x 200 mm resulting in an average voxel size of 11.25 ml. The grid was oriented to place voxels in the interventricular septum. Two saturation bands were placed over the skeletal muscle in the chest wall and one over the liver to minimise signal contamination.

The acquisition was non-gated with TR around 910-1010 ms depending upon SAR. The acquisition delay was reduced to a minimum (TE* = 0.3 ms) using the ultra-short echo time (UTE-CSI) technique to maximise acquired signal and reduce first order phase effects (therefore reducing artefact). The optimised RF pulse (duration 2.4 ms) was centred between γ and α peaks (usually by subtracting 250 Hz from the observed phosphocreatine frequency) to ensure uniform excitation^58^. Exploitation of the Nuclear Overhauser effect (NOE) was used to increase the signal-to-noise ratio in acquired spectra: five pulses, length 2.5 ms, inter-pulse delay 80.5 ms, pulse voltage 222.5 V and average flip angle 150°.

#### Spectral Analysis

The basal septal voxel was selected for analysis: this was the only user-dependent part of the process, the rest being fully automated. In-house Matlab software (OXSA^26^) determined the flip angle for the selected voxel by co-registering the short axis images with the spectral data. Flip angle variation due to coil loading effects was calculated using acquired inversion recovery data and the localizer containing the locations of cod liver oil capsules in fixed positions in the coil.

Pre-processing (baseline correction) was undertaken before fitting spectral peaks using the AMARES (advanced method of accurate, robust and efficient spectroscopic fitting) method. Peaks for phosphocreatine, α, β, γ - ATP, 2,3-diphosphoglycerate and phosphodiesters were fitted using prior knowledge of relative peak frequencies, J-coupling constants for ATP, relative peak amplitudes, relative phases and assumed Lorentzian line shapes along with acquisition parameters (central frequency, bandwidth, TR and calculated flip angles at various depths). Peak areas were corrected for RF partial saturation effects using the recorded excitation flip angle, T_1_ values (PCr 3.8s, γ -ATP 2.4s, α-ATP 2.5s, β-ATP 2.7s, 2,3-DPG 1.39s, PDE 1.11s) and spectral overlap with the NADH peak. The value of the ATP peak was corrected for blood contamination by subtracting 11% of the DPG peak area^58^.

PCr/ATP was calculated using the average of the three ATP peaks. The quality of spectral fit was assessed using the coefficient of variation in the measured PCr/ATP ratio, based on Cramer-Rao lower bounds (an indicator of signal to noise ratio in the sample) and standard error propagation formulae. Samples with a greater than 30% coefficient of variation were excluded.

### 4 Plasma Metabolomics and Proteomics Preparation and Analysis

#### Metabolomic profiling

Raw metabolomic data were processed using TraceFinder 3.1 (Thermo Fisher Scientific; Waltham, MA) and Progenesis QI (Nonlinear Dynamics; Newcastle upon Tyne, United Kingdom). LC-MS data were analyzed with Agilent Masshunter QQQ Quantitative analysis software. Isotope-labeled internal standards were monitored in each sample to ensure adequate MS sensitivity for quality control. Peaks were manually reviewed in a blinded fashion to assess quality.

#### Proteomic profiling

A Proximity Extension Assay technology was used, utilising pairs of oligonucleotide-labeled antibody probes to bind to their target protein. When the two probes are brought in close proximity, the oligonucleotides hybridise in a pair-wise manner. The addition of a deoxyribonucleic acid (DNA) polymerase results in proximity-dependent DNA-polymerisation, which creates a unique double-stranded DNA barcode for each specific antigen. Next-generation sequencing (Illumina NovaSeq) was then used to detect and quantify this DNA-sequence. Generated Data are quality-controlled and normalised using an internal extension control. The final assay readout is displayed as normalised protein eXpression (NPX) values, which are log2-transformed ratios of sample assay counts to extension control counts. All assay validation data are available on the manufacturer’s website (www.olinkexplore.com).

